# Illuminating histidine phosphorylation in the pancreatic tumor microenvironment

**DOI:** 10.1101/2022.09.15.508158

**Authors:** Natalie Luhtala, Nikki Lytle, Kathleen E. DelGiorno, Yu Shi, Razia Naeem, Michael A. Hollingsworth, Susan M. Kaech, Geoffrey M. Wahl, Tony Hunter

## Abstract

Development of phosphohistidine (pHis) antibodies has significantly advanced our understanding of pHis contributions to tumor biology, including a tumor suppressive role for a pHis phosphatase, a metastasis suppressive role for His kinases, and pHis regulation of T cell receptor signaling. Using these antibodies, we investigated pHis pathway regulation in the mouse pancreatic tumor microenvironment. We identified deregulated expression of pHis and pHis phosphatases that correlated with mouse pancreatic tumor progression. We developed a protocol to circumvent the acid and heat-sensitivity of pHis signals, enabling their co-staining with other proteins in FFPE tissue, identifying a significant enrichment of 1-pHis and a subtype of 3-pHis signals (Gly-3-pHis) in the stroma. We discovered increased Gly-3-pHis levels in tumor-associated myeloid cells mainly resulting from elevated ATP citrate lyase 3-pHis levels and predicted the existence of pHis in cell-cell adhesion proteins. We provide evidence that mitochondrial delocalization of PGAM5, a pHis phosphatase with increased expression during pancreatic tumorigenesis, occurs in tumor cells as compared to stromal cells, enabling access to PGAM5’s known cytoplasmic substrate, pHis-NME (Non-MEtastatic), and two potential Gly-3-pHis substrates, SCSα (Succinyl CoA Synthetase) and β-catenin. Overall, we introduce a new method and possible targets for future studies of pHis pathway deregulation during tumorigenesis.

## Introduction

More than 20 years ago, the discovery of phosphotyrosine (pTyr) revolutionized cancer research and treatment (Cohen et al., 2021; Huang et al., 2020; Hunter, 2015; Lipsick, 2019). Although histidine phosphorylation of Succinyl CoA Synthetase (SCS), a mitochondrial enzyme, was reported over 40 years earlier than pTyr(Boyer et al., 1962), histidine phosphorylation, an acid- and heat-sensitive modification, has more recently emerged as an area of interest in cancer biology, due to the development of isomer-specific anti-phosphohistidine monoclonal antibodies (Fuhs & Hunter, 2017; Hunter, 2022).

Individual phosphohistidine (pHis) regulators (names, pHis functions, localization, substrates, and relevance to cancer are detailed in Table 1) have been characterized for their functions in promoting or inhibiting tumorigenesis, invasion, and metastasis. NME1 (nucleoside diphosphate kinase 1), is a His kinase that generates pHis in proteins and whose function is linked to metastasis suppression(Khan & Steeg, 2018). LHPP (phospholysine, phosphohistidine inorganic pyrophosphate phosphatase), is a putative pHis phosphatase identified as a tumor suppressor(Hindupur et al., 2018). These proteins exhibit varying levels of expression and enhancement or inhibition of tumorigenesis and metastasis in different cancers(Adam, Lesperance, et al., 2020; Adam, Ning, et al., 2020; Tan & Chang, 2018).

**Table 1.**
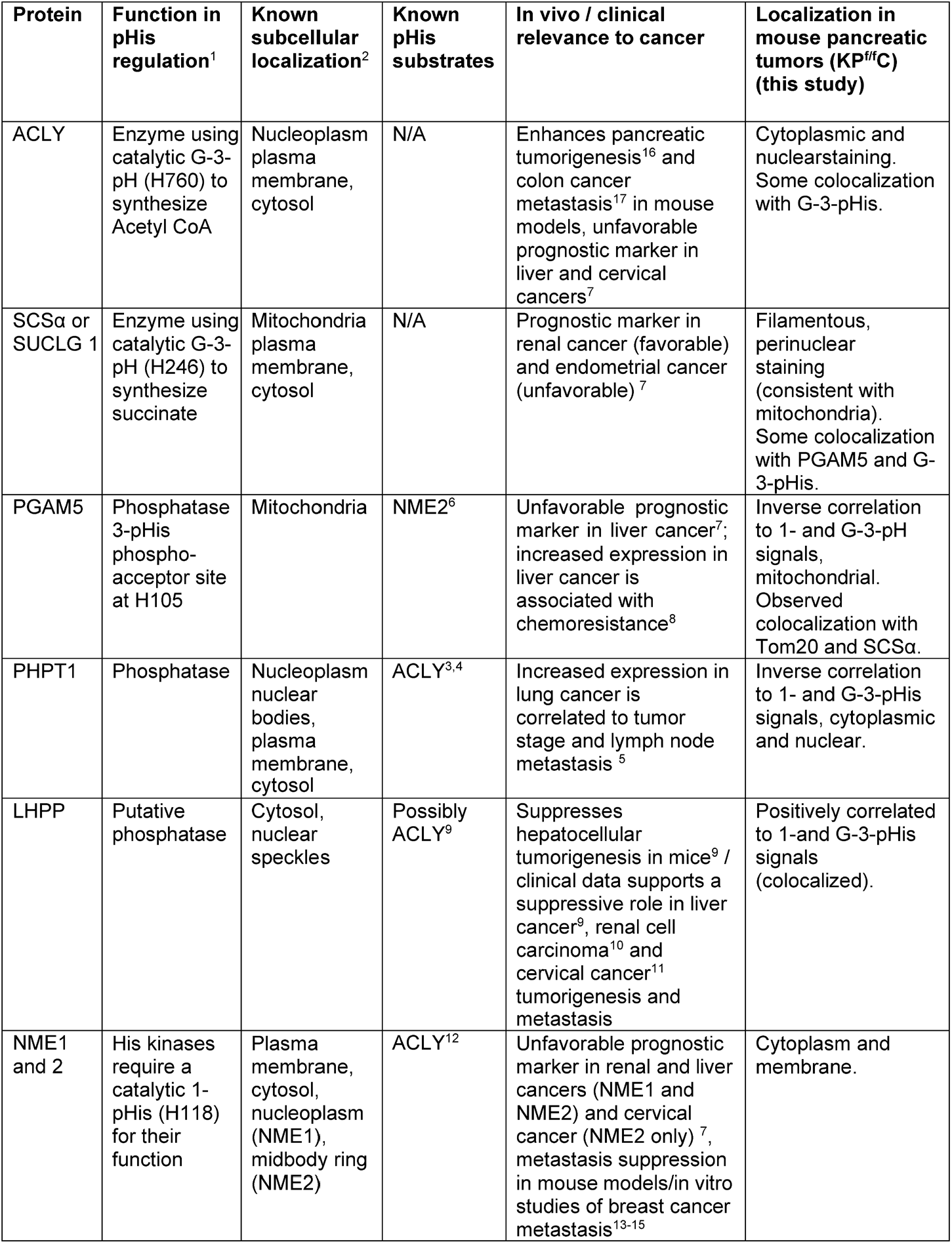
Phosphohistidine (pHis) proteins and regulators studied.

Other His kinases and pHis phosphatases exhibit multiple layers of reversible pHis regulation to regulate the opening and closing of T cell ion channels. The NME2 (nucleoside diphosphate kinase 2) His kinase positively regulates pHis on KCa3.1 (intermediate-conductance Ca^2+-^activated potassium channel 4), opening this channel. In contrast, protein histidine phosphatase 1 (PHPT1) dephosphorylates pHis on KCa3.1 and closes the channel. The PGAM5 (phosphoglycerate mutase family member 5) pHis phosphatase dephosphorylates 1-pHis in NME2 kinase and inactivates its function(Panda et al., 2016; Srivastava et al., 2006; Srivastava et al., 2008), preventing NME2’s His kinase activity against KCa3.1. Thus, examining the expression and localization of pHis signals and their regulators in a biologically appropriate context is needed to assess pHis signaling states in different cancer backgrounds and other models for disease. Such information could aid in defining therapeutic strategies tailored to pHis status in the tumor microenvironment.

Pancreatic ductal adenocarcinoma (PDAC) carries an abysmal prognosis with a 5-year survival rate of only 11 percent, shrinking to 3 percent for patients with regional metastases. Although a role for Acetyl CoA and the pHis enzyme ATP citrate lyase (ACLY) in PDAC tumorigenesis has been established (Carrer et al., 2019), a characterization of pHis regulation in PDAC tumorigenesis and metastasis is lacking. Both tumor cell intrinsic and extrinsic (cancer-associated fibroblasts, immune cells) signaling factors contribute to PDAC therapeutic resistance. We aimed to develop a protocol to holistically examine pHis regulators (kinases, phosphatases) and pHis signal expression in the three-dimensional pancreatic tumor microenvironment. To overcome the chemical instability of pHis antigens to traditional phosphoprotein analysis, we developed 1- and 3-phosphohistidine (pHis) isomer-specific rabbit monoclonal antibodies (mAbs)(Fuhs et al., 2015), enabling multiple types of pHis analyses, including immunoblotting, immunocytochemistry, and immunohistochemistry (IHC) staining of single markers in fixed, frozen tissue(Adam & Hunter, 2020; Kalagiri et al., 2020; Luhtala & Hunter, 2020). To better understand the spatial aspects of pHis signal regulation by His kinases and pHis phosphatases in tumors and to localize pHis signals to specific proteins, we developed an immunolocalization protocol for multiplexing pHis signals with those of pHis regulators and pHis proteins. Acid and heat treatment of pHis proteins significantly decreases or ablates their pHis signals, limiting multiplexed detection of pHis signals with other proteins whose detection requires traditional antigen retrieval methods. To circumvent this problem, we investigated the use of tyramide signal amplification (TSA)(Faget & Hnasko, 2015) to covalently attach fluorophores near the pHis antigen site in formalin-fixed paraffin-embedded (FFPE) mouse PDAC samples which can be multiplexed with pHis regulators and other marker proteins for spatial analyses.

Here, we developed an IHC-IF-pHis-TSA protocol to co-stain pHis signals with pHis regulators, pHis substrates, and cell type-specific markers in mouse PDAC FFPE sections. We characterized 3-pHis signals using the sequence-specific (GpHAG) anti-3-pHis mAb, SC44-1. The structural basis of SC44-1 mAb binding to 3-pHis-containing antigens has recently been characterized showing that it prefers a GpHAG motif(Kalagiri et al., 2021), enabling it to efficiently detect a catalytic Gly-3-pHis (G-3-pHis) on two important metabolic enzymes expressed in pancreatic tumors: ACLY, which enhances pancreatic tumorigenesis in mouse models(Carrer et al., 2019), localizing to the nucleus and the cytosol to catalyze the synthesis of acetyl CoA,(Fan et al., 2012; Sivanand et al., 2017) and succinyl CoA synthetase alpha subunit (SCSα), which in complex with SCSβ couples the synthesis of succinate from succinyl CoA in the mitochondria with the formation of ATP or GTP, depending on the organism(Huang & Fraser, 2021).

We co-stained for 1-pHis or G-3-pHis signals in mouse PDAC tumors with markers for tumor and stroma, and with signals for total pHis enzymes or three pHis phosphatases whose protein expression was increased in mouse PDAC tumors as compared to normal pancreatic tissue. Our results predicted the presence of histidine phosphorylated cell-cell adhesion proteins and unexpected ACLY G-3-pHis expression in PDAC tumor-associated myeloid cells (TAMs) and flagged PGAM5 mitochondrial phosphatase as a possible pHis regulator and therapeutic target during PDAC tumorigenesis and metastasis.

## Results

### Deregulated expression of pHis pathway proteins during pancreatic tumorigenesis

To understand the role of pHis regulation during the transition from normal pancreas to PDAC tumor, we performed immunoblots for pHis and pHis regulators (Table 1) of tissue lysates from mouse PDAC tumors from three different genetic models (*Kras*^*LSL-G12D/+*^; *p53*^*f/f*^; *Pdx1-Cre* = KP^f/f^C, *Kras*^*LSL-G12D/+*^; *p53*^*f/+*^; *Ptf1a-Cre* = KP^f/+^C, and *Kras* ^*LSL-G12D/+*^; *Ptf1a-Cre* = KC) as compared to normal pancreatic tissue (*Kras*^*LSL-G12D/+*^ = K and *p53*^*f/f*^; *Pdx1-Cre* = P^f/f^C), using normalized signals. We immunoblotted equal protein amounts of total tissue lysates using two anti-pHis mAbs: SC44-1 anti-3-pHis mAb (recognizes G-3-pH-AG found on ACLY and SCSα, hereafter referred to as anti-G-3-pH) and SC1-1 anti-1-pHis mAb (sequence-independent mAb that primarily recognizes NME1/2 1-pHis, hereafter referred to as anti-1-pHis). We also probed overall protein expression of enzymes phosphorylated at their catalytic His residue (pHis enzymes, ACLY and SCSα), kinases (NME1/2), and phosphatases (PHPT1, PGAM5, LHPP). Normalized signals were calculated (Table 2), and we examined target protein expression in mouse PDAC tumors compared to normal pancreatic tissue. One should bear in mind that the cellular compositions of normal pancreata and pancreatic tumor tissue are very different, and this needs to be borne in mind when interpreting any differences observed in the levels of proteins and protein modifications in whole tissue extracts, which requires detailed analysis of the distinct cell types in each tissue (see below).

**Table 2.**
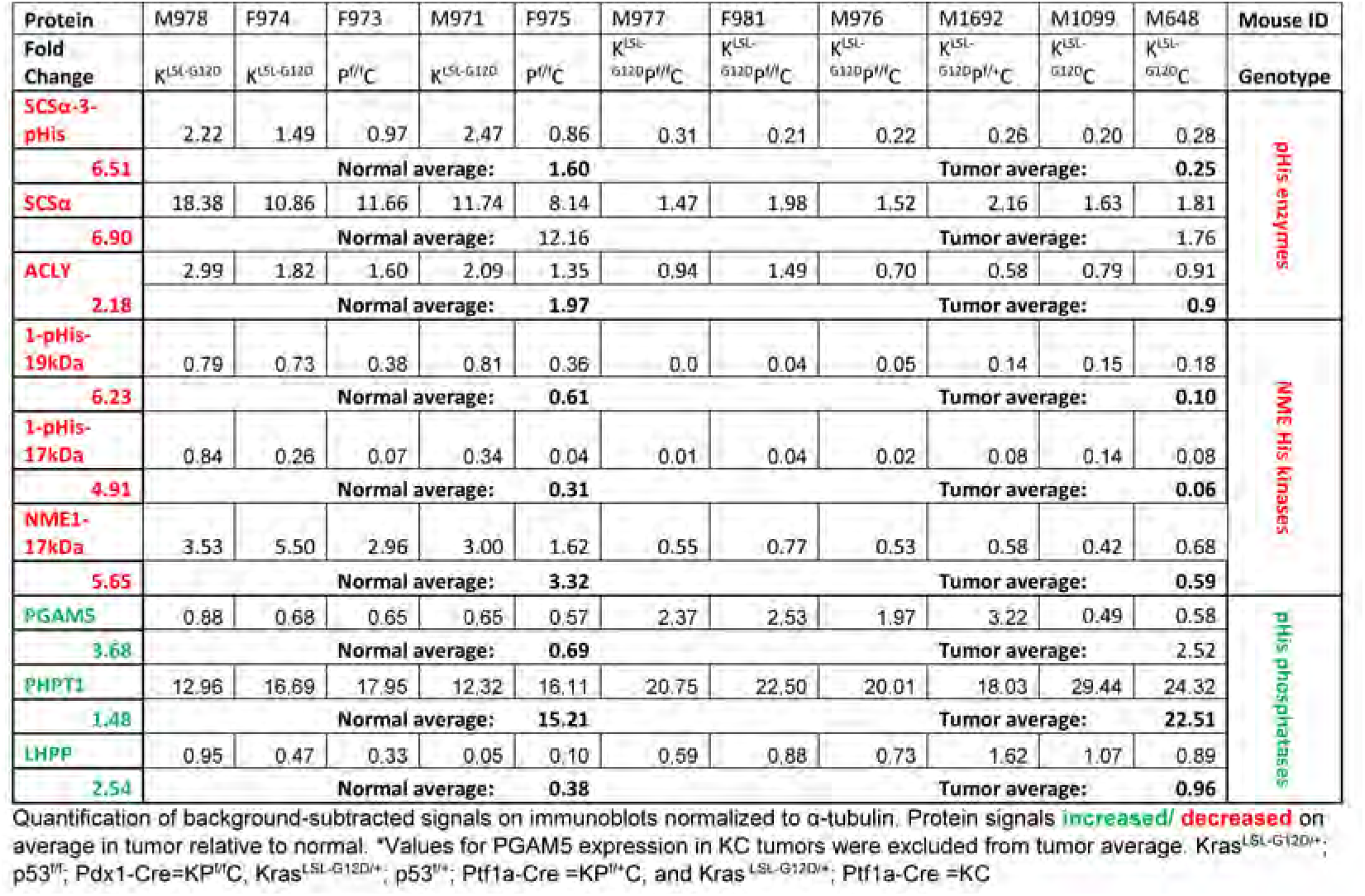
Normalized protein expression in mouse normal/tumor pancreatic tissue lysates.

We found that levels of both isoforms of pHis were decreased in mouse PDAC tumors, whereas expression of all tested pHis phosphatases was increased (Table 2). SCSα G-3-pH was detected more strongly in normal pancreas and was significantly reduced as compared to tumors (Fig 1A, E, 6.5-fold, p=0.004), and 1-pHis signals (likely NME1 or 2) demonstrated a similar pattern of decrease (6.2-fold, p=0.004 for the 19 kDa 1-pHis form, Fig 1B, E). Levels of three pHis phosphatases, PGAM5 (3.7-fold, p=0.016), PHPT1 (1.5-fold, p=0.004), and LHPP (2.5-fold, p=0.05) were significantly increased in PDAC tumors as compared to normal pancreas (Fig 1A, C, D, G).

**Fig. 1:**
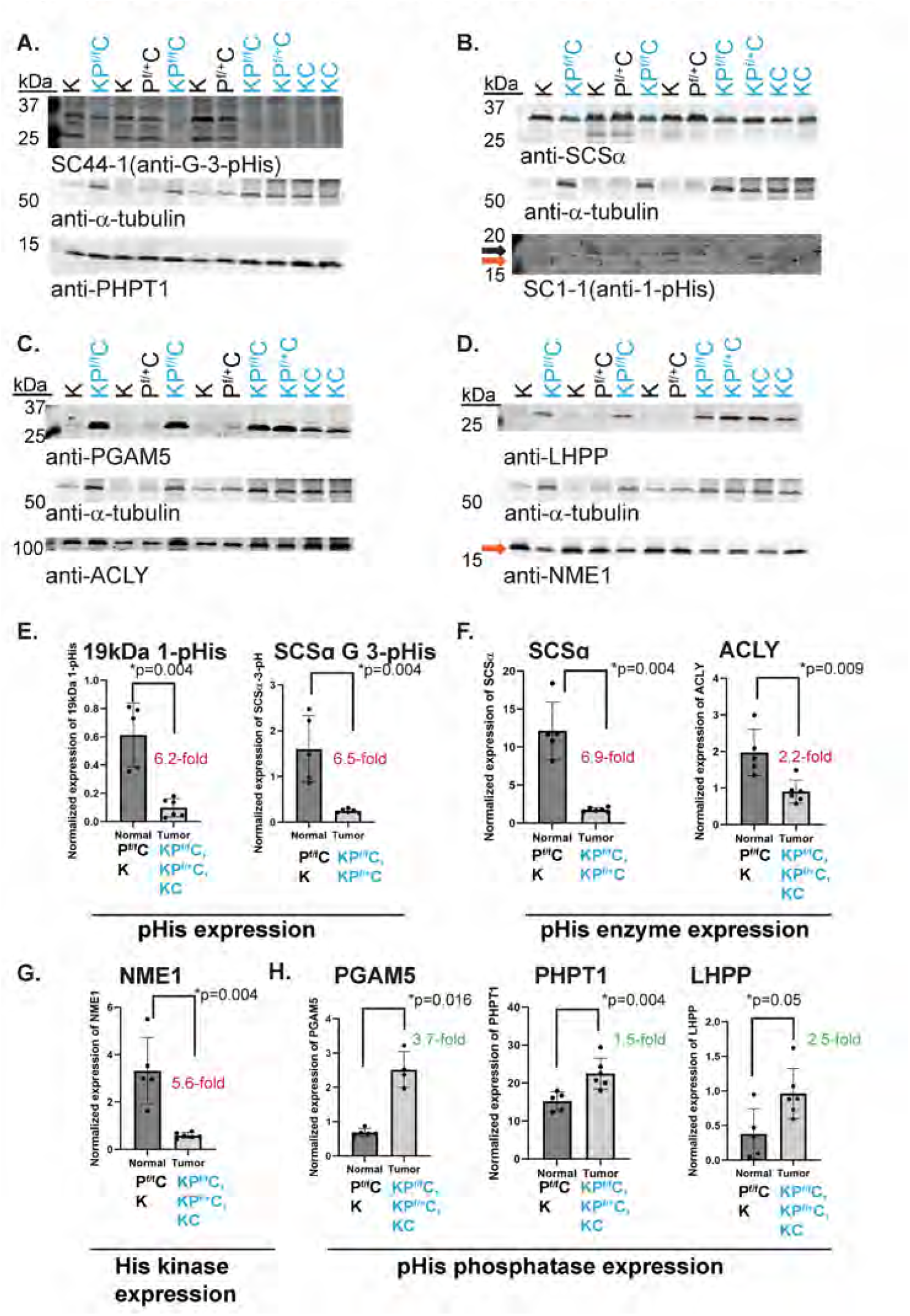
Altered expression of pHis and pHis regulators in mouse pancreatic ductal adenocarcinoma (PDAC) tumors. Immunoblotting (IB) and quantitation of tissue lysates from mouse normal pancreatic tissue=black font (K, P^f/f^C) or PDAC tumors=blue font (KC, KP^f/f^C, KP^f/+^C). (**a-d**) Immunoblots (IB) of protein expression are shown, each representing a single IB probed serially. Quantitation of IB (**e-h)** is shown as a graph of α-tubulin normalized expression (Mann-Whitney). Arrows indicate NME1 and likely NME pHis signals 17 kDa (orange) 19 kDa (black). Average fold increase (green) and decrease (red) is noted. pHis=phosphohistidine. FVB background: K, P^f/f^C, and KP^f/f^C=Kras^LSL-G12Df/+^; Tp53^f/f^; Pdx1-Cre. C57BL/6 background: KC and KP^f/+^C=Kras^LSL-G12Df/+^; Tp53^f/+^; Ptf1a-Cre.

Concordant with pHis signals, total pHis protein levels were decreased in mouse PDAC tumors relative to normal pancreas. By comparison, the expression of total NME1 protein was reduced (Fig 1D, G, 5.6-fold, p=0.004) as was the expression of total SCSα (Fig 1B, F, 6.9-fold, p=0.004). Of note, the decreased level of total and G-3-pH SCSα in tumors reached significance only in analyses of tumors lacking at least one copy of *TP53*, suggesting that either deregulation of p53 expression or a more advanced tumorigenic state is required for this transition (Fig 1E-F, Table 2). Although the ACLY G-3-pH signal was obscured by background on these immunoblots and was not quantitated, expression of total ACLY, previously shown to be important for mouse PDAC tumorigenesis(Carrer et al., 2019), was decreased in PDAC when normalized to α-tubulin (Fig 1C, F, 2.2-fold, p=0.009).

Increased expression of multiple pHis phosphatases and decreased levels of pHis protein signals (SCSα and NME) in mouse PDAC tumors as compared to normal pancreas introduces the possibility that dephosphorylation of pHis substrates occurs during pancreatic tumorigenesis. To correlate pHis proteins and phosphatases with pHis signals mapped spatially within the tumor microenvironment, we developed a protocol that circumvents the acid and heat sensitivity of pHis signals, enabling their co-staining with other proteins of interest.

### Conventional IHC on mouse pancreatic FFPE sections reveals differential localization of G-3-pHis signals

We aimed to develop a multiplex immunofluorescence staining protocol for co-localizing pHis signals and pHis regulators in normal mouse pancreas and advanced stage KP^f/f^C PDAC tumors. As a first step, we prepared sections of formalin-fixed paraffin-embedded (FFPE) tissue, using a modified, shorter FFPE protocol to preserve pHis signals, from 40–45-day-old littermates and tested whether pHis antigen signals could be detected using our antibodies in FFPE sections without antigen retrieval (AR) to avoid using acid and/or heat.

Following block preparation and sectioning, we probed for anti-G-3-pHis signals using the SC44-1 mAb and an HRP-conjugated secondary antibody combined with DAB staining, and hematoxylin counterstain (Fig 2A). Serial sections stained for defined tumor regions (anti-cytokeratin-19, CK19) and for one type of cancer-associated fibroblasts (CAFs) within the stroma (anti-alpha smooth muscle actin, αSMA) (Fig 2A, D, E), or with hematoxylin and eosin (H&E) staining were included in the analysis. Pre-blocking of the SC44-1 anti-G-3-pHis antibody with a His peptide phosphorylated by phosphoramidate (pept+PA) was used as a negative control, as validated previously(Luhtala & Hunter, 2020).

**Fig. 2:**
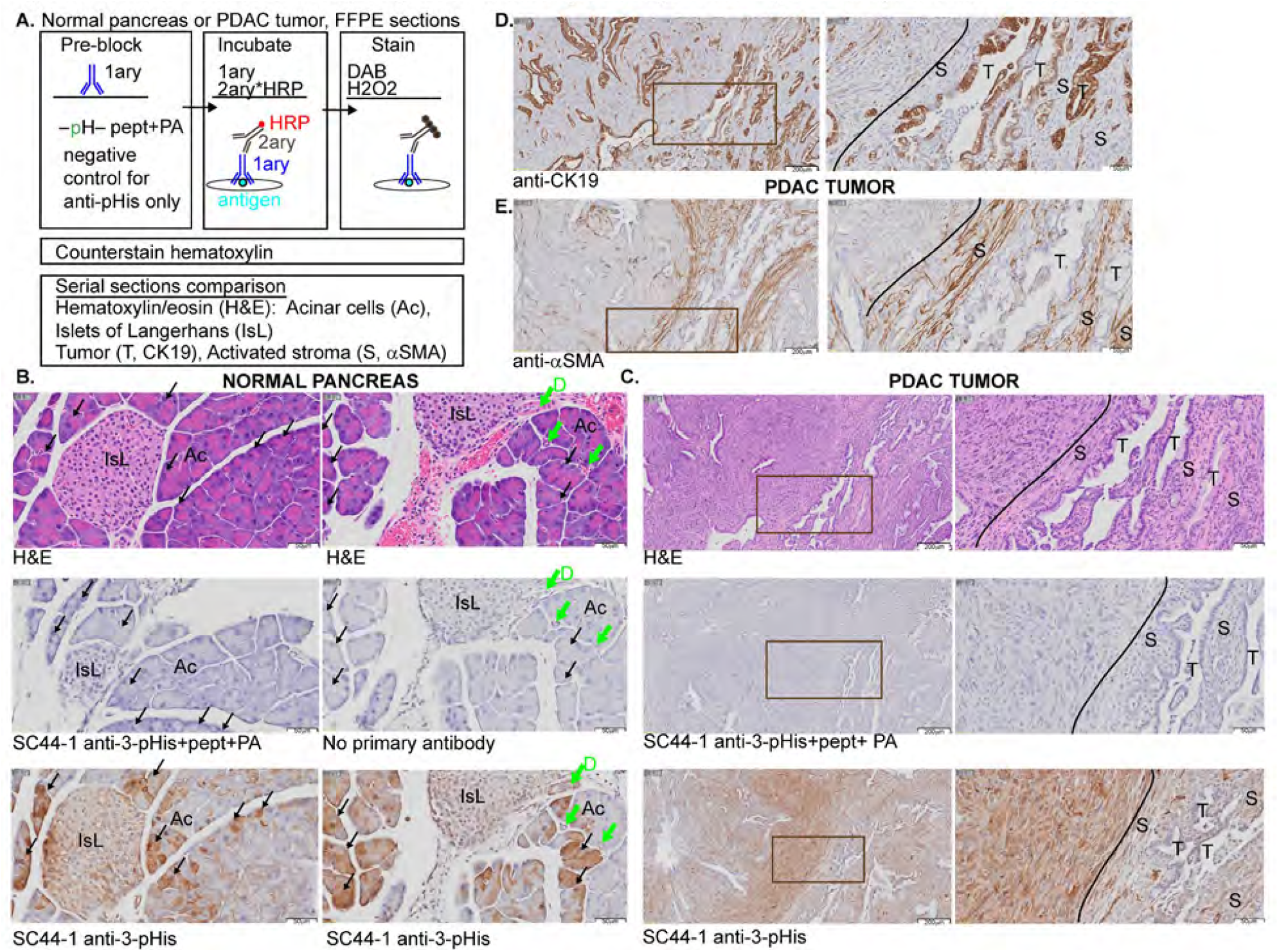
Differential localization of SC44-1 anti-G-3-pHis signals is revealed by chromogenic staining of mouse pancreatic FFPE sections. (**a)**, Schematic depicts the protocol used in this analysis. (**b-e)**, Chromogenic or H&E staining of pancreatic FFPE serial sections probed with the indicated antibodies. Negative control pept+PA indicates pre-blocking of SC44-1 G-3-pHis mAb prior to staining with phosphohistidine peptide. (**b)**, Normal mouse pancreas: black arrows indicate acini (Ac) and provide examples of strong staining. IsL labels islets of Langerhans; green arrows indicate ductal (D) regions. (**c-e**), Mouse PDAC tumors: the regions enclosed in outlined boxes in the left-hand panels are shown at higher magnification in the right panels, tumor (T) and cancer-associated fibroblast or activated stroma (S) regions were deduced by serial sections analysis of CK19 and αSMA staining, respectively. A line indicates the boundary of strong staining for G-3-pHis signals. Scale bars are indicated at the lower right corner of images. pHis=phosphohistidine.

We observed differential, *bona fide* pHis staining of cells in normal pancreas and PDAC tumors. Pre-blocking of the anti-G-3-pHis antibody eliminated signals to the same degree as staining without primary antibody (Fig 2B-C). In normal pancreas, the strongest signals were observed in a subset of acinar cells (Fig 2B, ‘Ac’, black arrows), which can undergo acinar-to-ductal metaplasia (ADM), forming preneoplastic regions. Ductal cells (green arrows, ‘D’) and Islets of Langerhans (‘IsL’) also expressed enhanced signals in a small percentage of cells, albeit less prominently than in acinar cells (Fig 2B). Differential detection of G-3-pHis signals in ducts, islets, and acini of normal pancreas and PDAC tumor and stroma suggests that cell and tissue context could affect the types of G-3-pHis-positive proteins present or their regulation by His kinases or pHis phosphatases.

In KP^f/f^C PDAC tumor tissue, regions most distant from tumor cells (left of the line, Fig 2C-E) displayed the strongest G-3-pHis signals. Signals for G-3-pHis were more pronounced in αSMA+ CAFs within the stroma (αSMA, S) than in regions of CK19-positive ductal cells (T) (Fig 2C-E) as evidenced by serial sections analysis of both low-(left) and high-magnification (right) images.

### IHC-IF-pHis-TSA protocol detects bona fide pHis signals in mouse PDAC FFPE sections

Signals for pHis are acid- and heat-sensitive, which precludes co-staining for pHis and other proteins and tumor/stromal markers whose staining requires antigen retrieval (AR) within a single tissue section. Consequently, we investigated the use of tyramide signal amplification (TSA) to covalently attach Cy5 fluorophores to tyrosines near the antigen site(Adams, 1992; Bobrow et al., 1989; Gross & Sizer, 1959) (Fig S1A) prior to AR, which involves boiling for 10 min. The covalent tyramide signals generated by the TSA staining protocol after incubations with anti-pHis primary antibody and a secondary HRP-conjugated antibody are stable to the denaturing conditions used for AR. Therefore, the use of TSA should permit subsequent AR and co-staining for pHis and other proteins of interest, including pHis regulatory proteins and biological marker proteins, on these FFPE sections.

Using serially sectioned FFPE mouse KP^f/f^C advanced stage tumor tissue, Cy5 far-red fluorophore, and Hoechst 33342 nuclear stain, we compared sections stained with anti-G-3-pHis mAb alone or pre-incubated with unmodified peptide, phosphoramidate alone (PA), or peptide phosphorylated by phosphoramidate (pept+PA) to produce pHis to block antibody binding. Analysis of scanned slides demonstrated strong pHis-dependent signals, that were lost with a pept+PA block, but not a pept or PA alone block (Fig S1B). The blocking effect of pept+PA was stronger than that observed with AR treatment prior to anti-pHis staining (Fig S1B, AR).

To validate the ability of this protocol to detect signals for the other isoform, 1-pHis, a modification present primarily on the NME1 and NME2 His kinases, we used the SC1-1 anti-1-pHis mAb. Due to the low level of 1-pHis generated by phosphoramidate (PA)-mediated phosphorylation of peptides used for blocking, we limited our controls to the use of AR for these analyses. Strong signals for 1-pHis were absent when AR was carried out prior to staining; the more complete loss of 1-pHis signals with AR treatment compared with the more moderate loss of G-3-pHis signals after AR is consistent with the reported lower stability of 1-pHis(Hultquist et al., 1966) (Fig S1C).

### G-3-pHis signals are enriched in the stroma of mouse PDAC

To map anti-G-3-pHis signals within the tumor microenvironment, we co-stained FFPE sections from KP^f/f^C PDAC advanced stage tumors as in Fig 2. We used antibodies against CK19 (tumor), αSMA (CAFs), or CD45 (immune cells) following a modified IHC-IF-pHis-TSA method.

We performed staining for G-3-pHis as in Fig S1, adding an AR step following completion of pHis TSA Cy5 staining. We then probed tissue sections with primary CK19 and secondary antibodies to detect tumor cells and performed a TSA step to attach Cy3 fluorophore near the CK19 antigen site, finishing with Hoechst nuclear staining and mounting to produce dual staining (Cy3/Cy5) for pHis (Cy5) and CK19 (Cy3).

As indicated by conventional IHC staining of tumor sections (Fig 2C-E), stronger signals for G-3-pHis mapped to stromal cells (lacking CK19 staining) in 2D (20X, Olympus slide scanning, Fig 3A) and 3D z-stack images (Zeiss Airyscan with Imaris analysis, Fig 3B). Low levels of nuclear/perinuclear G-3-pHis signals were observed in some tumor cells (yellow arrowheads, Fig 3A). QuPath analysis of co-stained G-3-pHis+CK19+ cell populations in multiple tumors demonstrated significant enrichment (mean=84%) of G-3-pHis signals in stromal (CK19-) cells (Fig 3G, p=0.0001). QuPath quantification revealed overall enrichment of 1-pHis signals in CK19-cells (mean=82%, Fig S2A). However, rare CK19+ tumor cell clones displayed 1-pHis staining on the basolateral side of tumor ducts (Fig S2B).

**Fig. 3:**
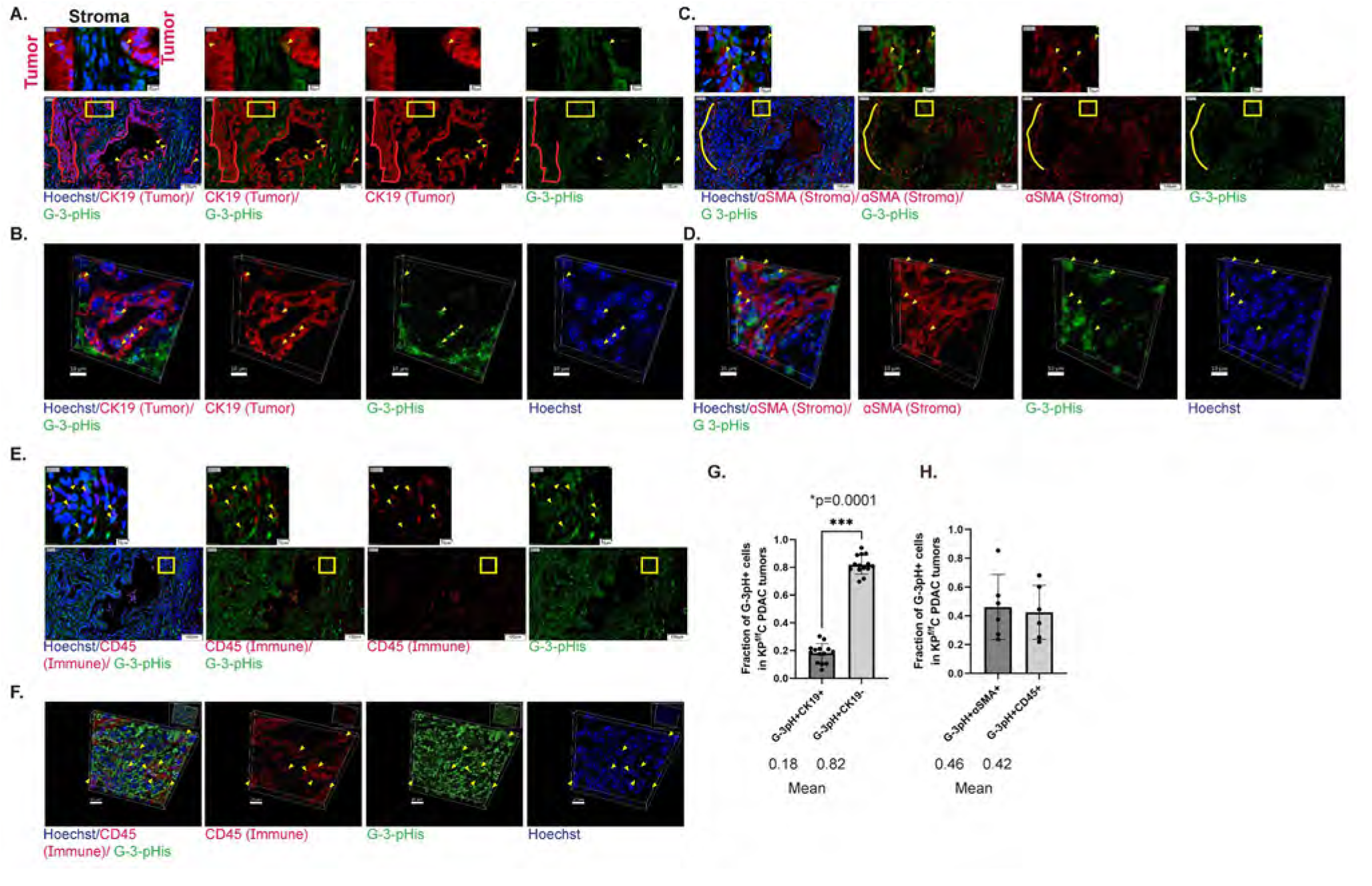
G-3-pHis signals are enriched in the stroma and immune cells of mouse PDAC tumors. (**a-f**) IHC-IF-pHis-TSA protocol (Fig. S1) applied to FFPE mouse KP^f/f^C PDAC tumor sections, using the indicated primary antibodies with TSA Cy5 (green signals, pHis) or TSA Cy3 (red signals, CK19, αSMA, or CD45) reagents, Hoechst staining of nuclei in blue. (**a, c, e**) Slide scanned images (20X, Olympus). The regions enclosed by the yellow boxes on the lower magnification images (bottom) are shown at higher magnification above. (**a**) Red outline denotes a CK19+ duct with little G-3-pHis staining (**c**) Yellow outline denotes G-3-pHis+ staining in stroma. QuPath quantitation of the fraction of G-3-pHis signals co-staining in KP^f/f^C tumors: (**g**) with CK19, n=12, ***p=0.0001, Wilcoxon) (**h**) with αSMA or CD45, n=6. (**b, d, f**) Super-resolution (Airyscan, Zeiss) z-series images rendered in 3D using Imaris. Yellow arrowheads show G-3-pHis signals in tumor (**a-b**), stroma (**c-d**), or immune cells (**e-f**). Scale bars are indicated in (**a, c, e**) at the lower right corner or in (**b, d, f**) at the lower left corner of images. pHis/pH=phosphohistidine, PDAC=pancreatic ductal adenocarcinoma.

To map G-3-pHis signals within two types of stromal cells, we co-stained for G-3-pHis signals and α-SMA+ CAFs (α-SMA, Fig 3C-D) or immune cells (cluster of differentiation 45 antigen, CD45, Fig 3E-F). CAFs (α-SMA+) and immune cells co-staining with G-3-pHis were readily detected in 2D and 3D images (yellow arrowheads, Fig 3C-F). QuPath analysis of G-3-pHis co-staining with α-SMA or CD45 in multiple KP^f/f^C tumors (Fig 3H) revealed that a greater fraction of G-3-pHis+ cells were either CAFs (α-SMA+, mean=46%) or immune cells (CD45+, mean=42%) rather than tumor cells (CK19+, mean=18%, Fig 3G).

### G-3-pHis proteomic analyses identify novel forms of ACLY and cell-cell adhesion proteins in mouse PDAC

To profile the proteins that correspond to G-3-pHis signals in PDAC, we dissociated KP^f/f^C PDAC advanced stage tumors and utilized fluorescence activated cell sorting (FACS) (profiled in Fig S3) to isolate tumor (EpCAM+), immune (CD45+;EpCAM-), and stromal (cancer-associated fibroblasts, CD45-;EpCAM-;CD31-;podoplanin+) populations. Using immunoblots of lysates from small numbers of cells (5×10^4^), we observed enrichment of G-3-pHis signals in immune cells. Specifically, we consistently observed a G-3-pHis+ 75 kDa protein (red arrow) in these cells not observed in other cell types (Fig 4A-B). Since the 120 kDa protein band (black arrow) we detected corresponded to the size of full-length ACLY, we used Odyssey IR scanner 2-color analysis (Fig 4C, total ACLY=red, G-3-pHis=green) and found that both the 120 kDa and 75 kDa protein bands and an intermediate-sized band observed for G-3-pHis immunoblots were also detected by anti-ACLY antibodies (ACLY G-3-pHis=yellow). As expected, G-3-pHis signals, but not total ACLY signals, were reduced by acid and heat treatment prior to SDS-PAGE and immunoblot (Fig 4C, note the presence of only total ACLY signals=red).

**Fig. 4:**
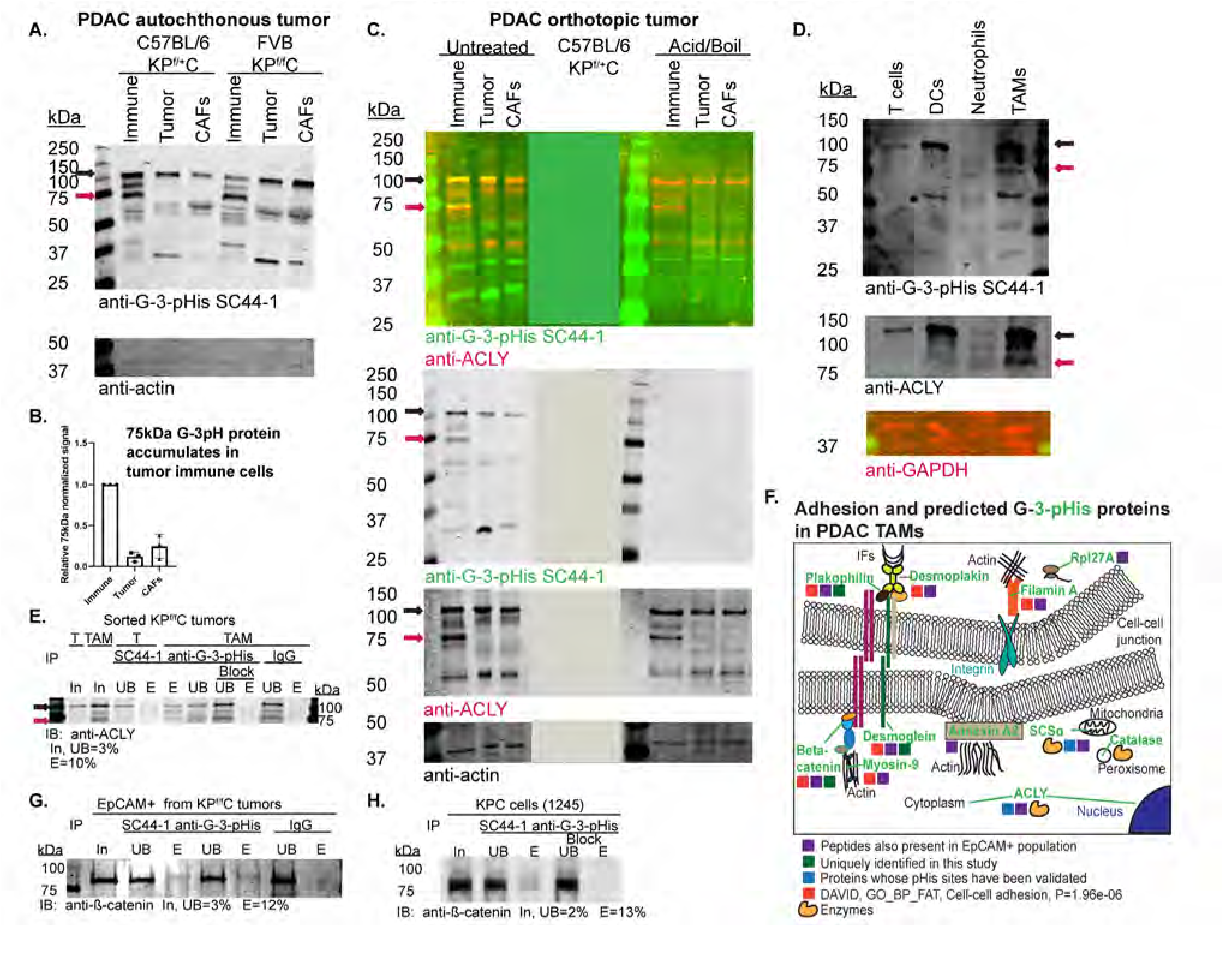
Mouse PDAC G-3-pHis proteomic analyses reveal cell-cell adhesion proteins and novel forms of ACLY. (**a, c, d**) Immunoblots of FACS populations obtained from the indicated mouse PDAC tumors; the position of canonical (120 kDa, black) and non-canonical (75 kDa, red) forms of ACLY G-3-pHis are indicated by arrows. Immune=CD45+EpCAM-, Tumor=EpCAM+, CAFs=EpCAM-CD45-CD31-Podoplanin+, T cells=CD3+, DCs=CD11c+ - contains certain dendritic cells, Acid/Boil=control to hydrolyze G-3-pHis signals, neutrophils=CD11b+Ly6G+, tumor-associated monocytes/macrophages (TAMs)=CD11b+Ly6G. (**b**) Quantitation of normalized immunoblot signals for the G-3-pHis 75 kDa band. (**e, g, h**) Immunoblots of fractions obtained from immunoprecipitation. In=input, UB=unbound, E=eluted, IgG=immunoglobulin control, block=SC44-1 incubation with 3-pTza peptide. (f) GH motif proteins identified by IP-SC44-1-MS that were enriched in TAMs as compared to IgG and peptide block controls (Supplemental File 1). pHis=phosphohistidine, PDAC=pancreatic ductal adenocarcinoma.

To identify which types of immune cells expressed the multiple unique G-3-pHis protein bands observed on immunoblots, we further sorted CD45+;EpCAM-cells from KP^f/f^C PDAC tumors into T cell (CD3+), dendritic cell (CD11c+), neutrophil (CD11b+;Ly6G+), and tumor-associated monocyte/macrophage (TAMs, CD11b+;Ly6G-) populations. TAMs were the most abundant immune population in tumors and showed clear expression of three G-3-pHis protein bands on immunoblots (Fig 4D).

To identify other G-3-pHis proteins in TAMs, we performed anti-G-3-pHis SC44-1 mAb immunoprecipitation/mass spectrometry (IP/MS) on EpCAM+ tumor cell and TAM populations from pooled advanced stage KP^f/f^C PDAC tumors, analyzing 3×10^5^ cells per IP. As negative controls, we performed IPs on fully denatured lysates from TAMs using rabbit IgG or SC-44-1 G-3-pHis antibody pre-incubated with a previously characterized 3pTzA peptide as a control(Fuhs et al., 2015). Prior to MS analysis, we validated the presence of ACLY in lysates (input, In) and eluates (E) from experimental IPs, but not negative controls (IgG, Block) (Fig 4E). Peptides predicted from MS spectra were analyzed, and predicted proteins were filtered based upon the presence of at least one GH peptide motif in the protein’s sequence and a greater number of peptides in the anti-G-3-pHis IP than negative controls (Supplemental File 1). GH motif proteins predicted for tumor and TAMs populations in this analysis showed some overlap with proteins previously identified in published pHis phosphoproteomic analyses (Adam, Lesperance, et al., 2020; Fuhs et al., 2015; Hindupur et al., 2018), and GH motifs were conserved from mouse to human (Supplemental File 1). Our analysis is the first to predict the presence of plakophilin, desmogleins, and β-catenin in anti-G-3-pHis immunoprecipitates, and functional annotation of the putative G-3-pHis proteome of TAMs revealed a significant enrichment for the biological process of cell-cell adhesion (DAVID, GO_BP_FAT, GO: 0098609, P=1.96e-06) (Supplemental File 1, Fig 4F). Plakophilin-1 (80.9 kDa) and catalase (59.8 kDa) were the only predicted G-3-pHis proteins identified in TAMs, but not tumor cells (Supplemental File 1).

We validated the IP/MS results for β-catenin in tumor cells, since a greater number of peptides were predicted by MS in IPs of tumor cells as compared to IPs of TAMs. As in Fig 4E, we analyzed total input (In), unbound (UB), and eluted (E) fractions from anti-G-3-pHis IPs from lysates of 3×10^5^ sorted KP^f/f^C tumor (EpCAM+) cells and from lysates of 4×10^6^ cultured 1245 KPC cells, established from a *Kras*^*G12D/+*^*;p53*^*R172H/+*^*;Pdx1-Cre* PDAC tumor(Boj et al., 2015). In both conditions, we observed that a small percentage of β-catenin was immunoprecipitated using anti-G-3-pHis mAb, but not in control IPs using rabbit IgG (Fig 4G) or anti-G-3-pHis mAb blocked with 3-pTza peptide (Fig 4H) as controls.

### G-3-pHis signals for ACLY or SCSα are less abundant on average in mouse PDAC tumor as compared to stroma

Since the most abundant peptides identified in our IP/MS assay represented ACLY and SCSα in both tumor and TAMs, we sought to understand which anti-G-3-pHis signals correspond to G-3-pHis ACLY or SCSα in the tumor and stroma of the pancreatic tumor microenvironment. To quantitate ACLY G-3-pHis and SCSα G-3-pHis signals in tumor and stroma at higher resolution, we performed triple staining of FFPE mouse PDAC tumor sections, co-staining for G-3-pHis (Cy5) and CK19 (Cy3) and ACLY or SCSα (fluorescein). Triple-stained z-series images were captured by super-resolution microscopy and quantitated using colocalization of ACLY (Fig 5A) or SCSα signals (Fig 5B) with G-3-pHis in CK19-masked (tumor, yellow surface) and unmasked (stroma) regions.

**Fig. 5:**
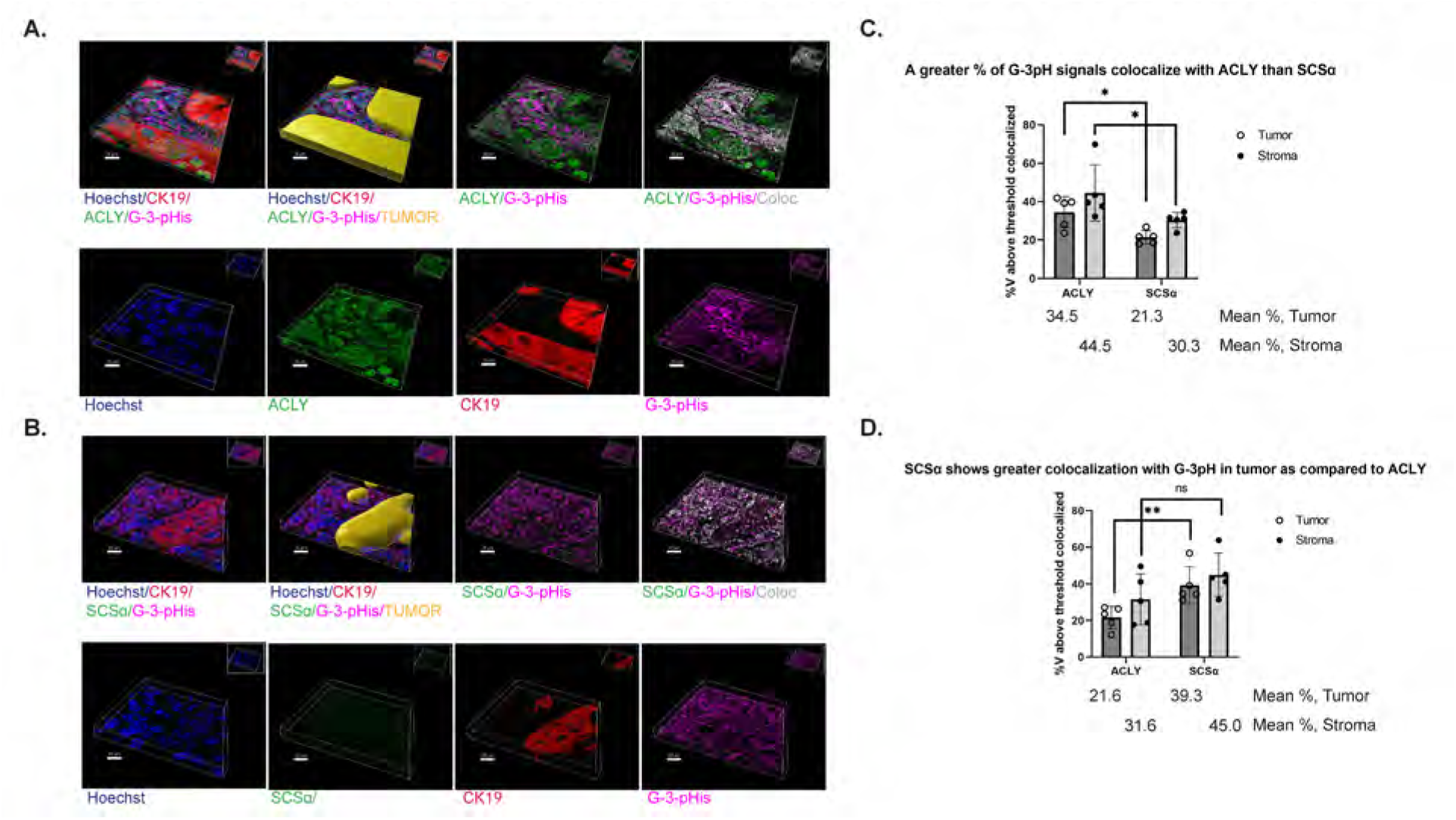
G-3-pHis signals for ACLY or SCSα are less abundant on average in mouse PDAC tumors as compared to stroma. (**a-b**), IHC-IF-pHis-TSA protocol (Fig S1) applied to FFPE KP^f/f^C mouse PDAC tumor using the indicated primary antibodies for triple-staining with TSA Cy5 (G-3-pHis, pink signals), TSA Cy3 (CK19, red signals), and TSA fluorescein (ACLY or SCSα, green signals)., Hoechst staining of nuclei in blue. Super-resolution (Airyscan, Zeiss) z-series images rendered in 3D using Imaris. Masked tumor is shown in yellow, colocalization is shown in black to white (low to high) to represent the intensity of colocalization. Scale bars are shown at the bottom left of images. Graphs of (**c**) %V of G-3-pHis signal colocalized with ACLY or SCSα in tumor or stroma (n=5, *p=0.0159, Mann-Whitney) (**d**) %V of ACLY or SCSα signal colocalized with G-3-pHis in tumor and stroma (n=5, **p=0.0079, Mann-Whitney) are shown. V=volume. pHis/pH=phosphohistidine.

Results indicated that ACLY G-3-pHis signals and SCSα G-3-pHis signals were slightly more abundant, albeit not significantly, in stroma as compared to tumor (Fig 5A-C). Colocalization of G-3-pHis signals with either pHis enzyme (Fig 5C) or pHis enzyme colocalization with G-3-pHis signals (Fig 5D) was slightly, but not significantly, decreased in tumors as compared to stroma. The percentage of G-3-pHis signal colocalization with ACLY was greater than that of SCSα in both tumor and stroma (Fig 5C). However, a significantly greater percentage of SCSα signals colocalized with G-3-pHis in the tumor, as compared to ACLY (Fig 5D). These data reveal that this IHC-IF-pHis-protocol can identify and localize specific histidine phosphorylated proteins in the tumor microenvironment.

### Signals for PHPT1 and PGAM5 histidine phosphatases are inversely correlated with pHis signals

In experiments comparing mouse PDAC tumor lysates to normal pancreatic tissue lysates, expression of the PGAM5, PHPT1, and LHPP pHis phosphatases was significantly increased (Fig 1M) while pHis levels (SCSα G-3-pH and 1-pHis, 19 kDa) were significantly decreased (Fig 1K). To explore these relationships further, we compared the localization of pHis phosphatases with pHis signals in the pancreatic tumor microenvironment, using IHC-IF-pHis-TSA analysis to co-stain FFPE sections from mouse KP^f/f^C PDAC tumors for PGAM5, PHPT1, or LHPP phosphatases (Cy3, red) and pHis signals (1-pHis or G-3-pH, Cy5, green), to investigate their colocalization and relationship with pHis signals in PDAC tumors and stroma. In interpreting such co-localization results, it should be stressed that at this level of resolution one cannot determine whether a pHis phosphatase molecule is close enough to a pHis substrate protein molecule to carry out direct dephosphorylation. Additional proximity-based studies would be needed to establish whether this is the case.

In scanned slide images (20X) we observed the strongest inverse correlation between G-3-pHis signals and PHPT1 pHis phosphatase signals. Even at low resolution, cellular regions exhibiting an inverse relationship in the strength of PHPT1 and G-3-pH signals (white outlines) could be identified (Fig 6A). Magnification of yellow boxed regions (above) reinforced this inverse relationship (white arrows, upper images, Fig 6A).

**Fig. 6:**
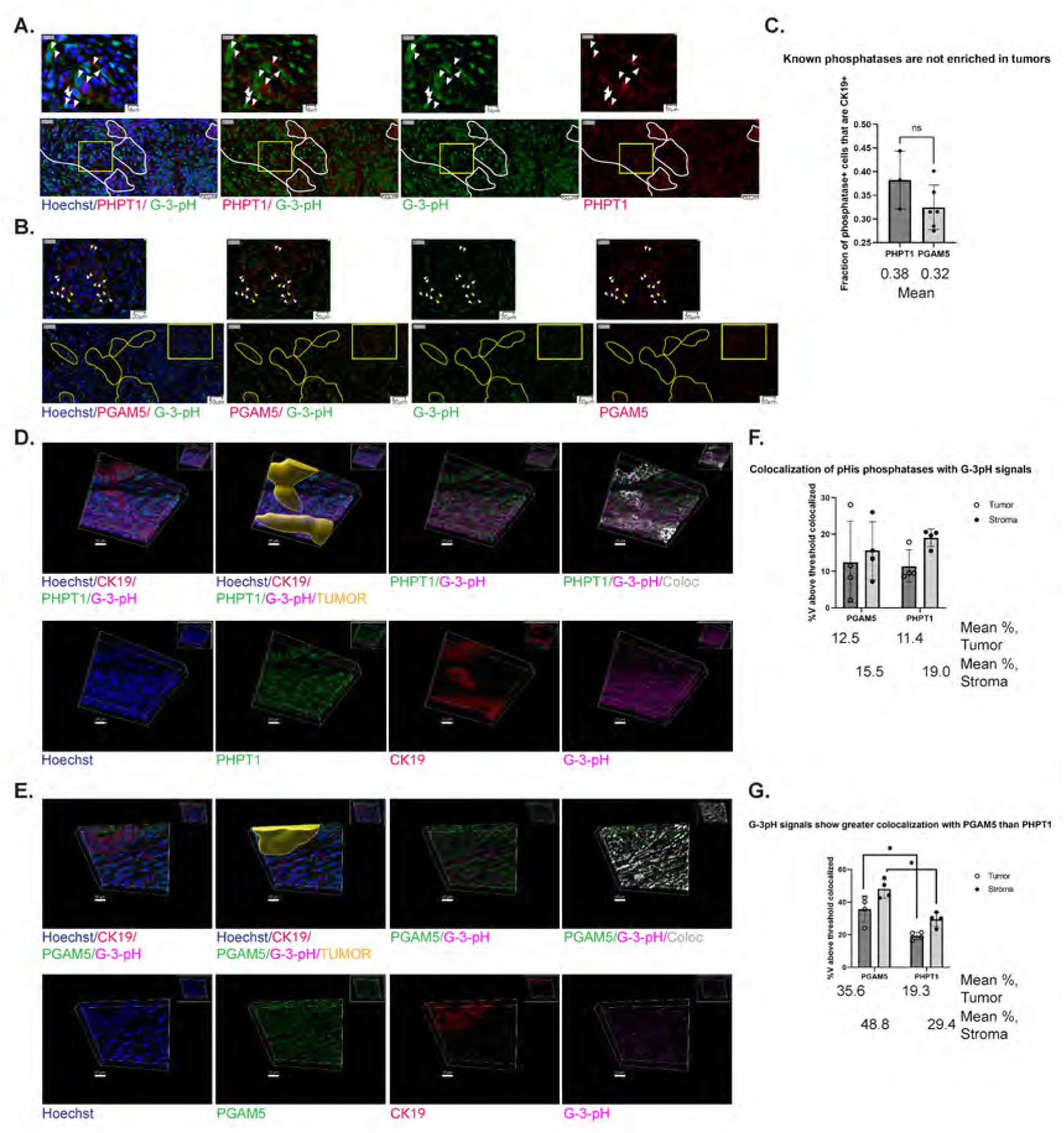
PHPT1 and PGAM5 signals exhibit an inverse relationship to pHis signals. (**a-b**), IHC-IF-pHis-TSA protocol (Fig S1) applied to FFPE KP^f/f^C mouse PDAC tumor serial sections analyzed using the indicated primary antibodies with TSA Cy5 (green signals, G-3-pHis) or TSA Cy3 (red signals, PHPT1 or PGAM5) reagents, Hoechst staining of nuclei in blue. Slide scanned images (20X, Olympus). The regions enclosed by the yellow boxes on the lower magnification images (bottom) are shown at higher magnification above. White outlines and arrowheads provide examples of red and green signals with an inverse relationship, yellow outlines and arrowheads display a less clear inverse relationship. Scale bars are indicated at the lower right corner of images. (**c**) QuPath quantitation of the fraction of G-3-pHis signals co-staining of PHPT1 or PGAM5 with CK19 in KP^f/f^C tumors. (**d-e**) As in (a-b), but triple-staining with TSA Cy5 (G-3-pHis, pink signals), TSA Cy3 (CK19, red signals), and TSA fluorescein (PHPT1 or PGAM5, green signals) was used. Super-resolution (Airyscan, Zeiss) z-series images rendered in 3D using Imaris. Masked tumor is shown in yellow, colocalization is shown in black to white (low to high) to represent the intensity of colocalization. Scale bars are shown at the bottom left of images. Graphs of (**f**) % of PHPT1 or PGAM5 signal colocalized with G-3-pHis signals in tumor or stroma (n=4) (**g**) % of G-3-pHis signal colocalized with PHPT1 or PGAM5 in tumor and stroma (n=4, *p=0.0286, Mann-Whitney) are shown. V=volume, pHis/pH=phosphohistidine.

By contrast, the relationship between PGAM5 and G-3-pHis signals was not as clear at low magnification where signals intermingled in multiple cellular regions (Fig 6B, yellow outlines). However, at higher magnification, we observed overlapping (yellow) and inversely related signals (white) (arrowheads, Fig 6B). Signals for 1-pHis were inversely related to both PHPT1 and PGAM5 signals (Fig S4, white lines and arrowheads), whereas LHPP demonstrated primarily colocalization with both 1-pHis and G-3-pH signals in 2D analyses (Fig S5, yellow arrowheads).

To map PHPT1 and PGAM5 signals with G-3-pHis signals at super-resolution in 3D, using CK19 to define tumor (CK19+) and stromal (CK19-) cells, we utilized triple co-staining as in Fig 5, staining for G-3-pHis (Cy5), CK19 (Cy3) and PHPT1 or PGAM5 (fluorescein). Scanned slide images were first analyzed using QuPath to determine the fraction of cells expressing PHPT1 or PGAM5 that are tumor cells (CK19+). In principle, enrichment of a pHis phosphatase in tumor cells could account for the decreased G-3-pHis signals observed in tumor as compared to stroma, but neither PHPT1 nor PGAM5 were enriched in tumor cells (Fig 6C).

Subsequently, we captured z-series images of triple-stained samples using super-resolution microscopy and quantitated colocalization of PHPT1 (Fig 6D) or PGAM5 signals (Fig 6E) with G-3-pHis in CK19-masked (tumor, yellow surface) and unmasked (stroma) regions. As suggested by QuPath 2D analysis (Fig 6C), signals for both pHis phosphatases colocalized with G-3-pH in cells of the tumor and stroma. However, a significantly greater percentage of G-3-pH signals colocalized with PGAM5 (in both tumor and stroma) than observed for PHPT1. Consistent with our observations in slide scanned images (Fig 6A-B), PHPT1 staining recorded stronger inverse Pearson’s correlation coefficients for the colocalized volumes than PGAM5 (Table 3), although, apart from one exception, correlation coefficients for pHis phosphatase signal colocalization with G-3-pH signals were all negative.

**Table 3.**
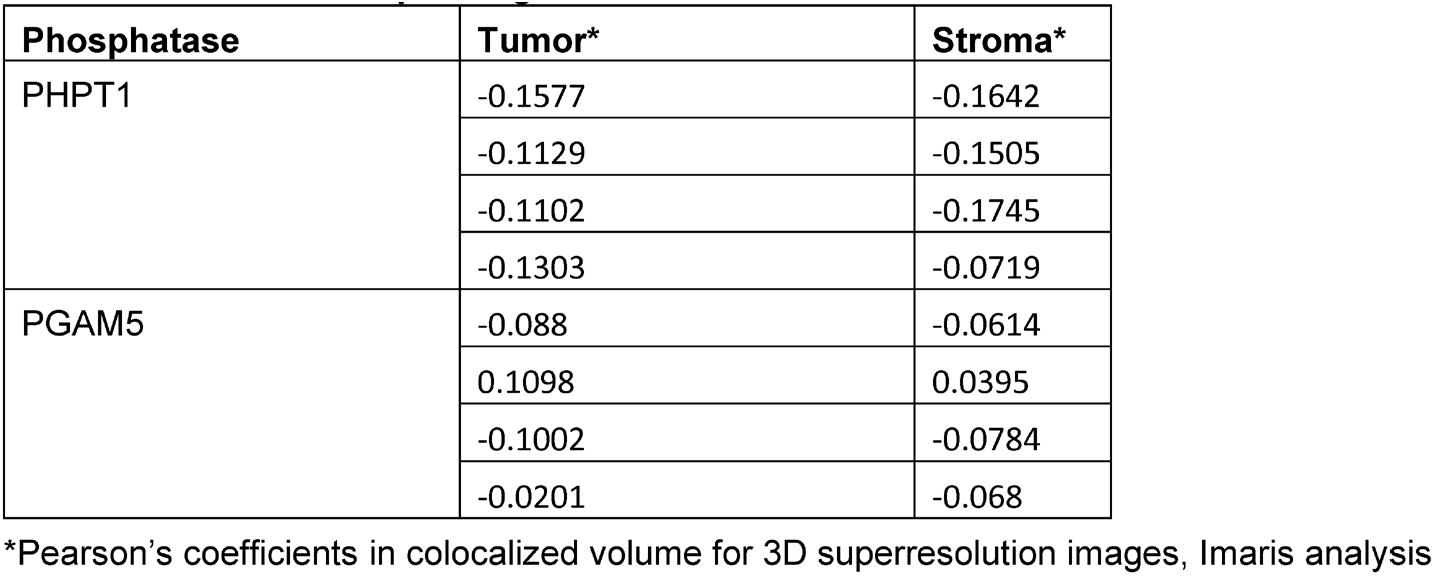
PHPT1 and PGAM5 phosphatases demonstrate inverse correlation coefficients in analyses of their colocalization with G-3-pHis signals.

### PGAM5 phosphatase colocalizes most with SCSα that is not associated with G-3-pHis in PDAC tumors

Next, we asked whether PGAM5, a mitochondrial phosphatase, might affect levels of G-3-pHis on SCSα, since both are mitochondrial proteins and the increase in PGAM5 protein expression in tumor tissue as compared to normal pancreas tissue was accompanied by decreased detection of SCSα G-3-pHis (Fig 1). To address this question, we triple-stained FFPE mouse PDAC KP^f/f^C advanced stage tumor tissue using IHC-IF-TSA and antibodies against G-3-pHis, SCSα, and PGAM5 and TSA staining to deposit Cy5, Cy3, and fluorescein fluorophores at each respective antigen site (Fig 7A). Using super-resolution images (z-series represented in 3D), we analyzed colocalization of PGAM5 with SCSα (Fig 7B, D) and compared this to G-3-pHis colocalization with SCSα (Fig 7C-D). Although on average, 24.4% of PGAM5 signals colocalized with SCSα, this was significantly less (p=0.0006) than observed for G-3-pHis colocalization with SCSα (46.8% mean, Fig 7D). Next, we compared PGAM5 signal colocalization with G-3-pHis signals to SCSα signal colocalization with G-3-pHis signals. We observed a significantly greater percentage (p=0.0012) of SCSα signals colocalized with G-3-pHis signals (Fig 7C,F, mean=30.9), as compared to the colocalization of PGAM5 signals with G-3-pHis signals (Fig 7E-F, mean=15.3).

**Fig. 7:**
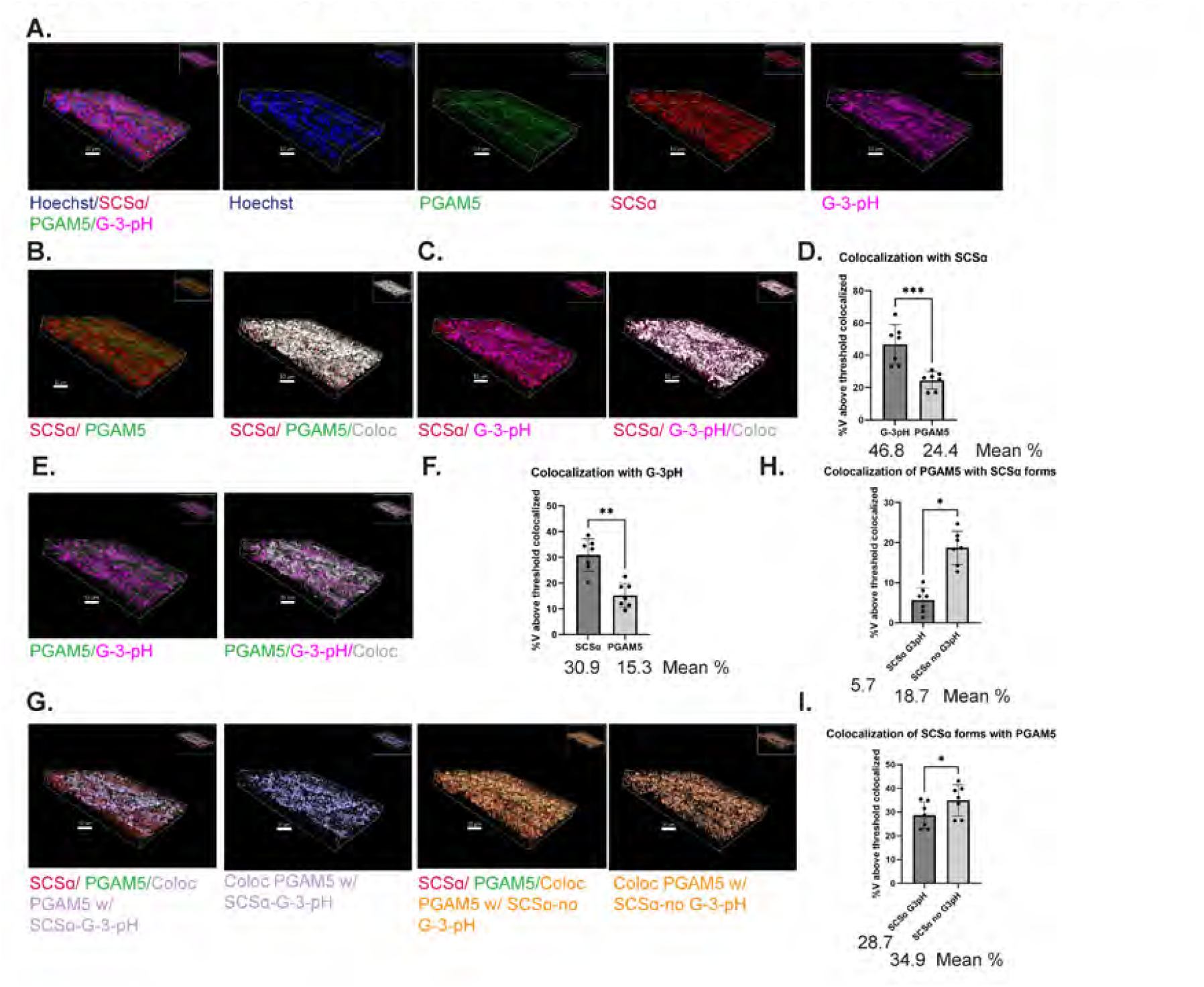
PGAM5 shows greater colocalization with SCSα that is not histidine phosphorylated. (**a, b, c, e, g**), IHC-IF-pHis-TSA protocol (Fig S1) applied to FFPE KP^f/f^C mouse PDAC tumor serial sections analyzed with triple-staining using TSA Cy5 (G-3-pHis, pink signals), TSA Cy3 (SCSα, red signals), and TSA fluorescein PGAM5, green signals). Super-resolution (Airyscan, Zeiss) z-series images rendered in 3D using Imaris. Scale bars are shown at the bottom left of images. (**a**) Merged and individual channel’s signals. Colocalization of SCSα with PGAM5 (**b**) or with G-3-pHis (**c**). (**d**) %V of G-3-pHis or PGAM5 colocalization with SCSα (n=7, ***p=0.0006, Mann-Whitney). (**e**) Colocalization of PGAM5 with G-3-pHis. (**b, c, e**) Colocalization shown in black to white (low to high) to represent the intensity of colocalization. (**f**) %V of SCSα or PGAM5 colocalized with G-3-pHis. (n=7, **p=0.0012, Mann-Whitney) (**g**) SCSα (red), PGAM5 (green), channel showing colocalization of PGAM5 with SCSα G-3-pHis (lilac), channel showing colocalization of PGAM5 with no G-3-pHis (orange). (**h**) %V of PGAM5 colocalized with SCSα G-3-pHis or SCSα not associated with G-3-pHis (n=7, *p=0.0156, Wilcoxon). With G-3-pHis. (**i**) %V of SCSα G-3-pHis or SCSα not associated with G-3-pHis colocalized with PGAM5 (n=7, *p=0.0469, Wilcoxon). Scale bars are shown at the bottom left of images. V=volume, pHis/pH=phosphohistidine.

These initial analyses demonstrated that a smaller percentage of PGAM5 signals was associated spatially with G-3-pH signals than was associated with SCSα protein, suggesting that PGAM5 preferentially co-localizes with SCSα lacking the G-3-pH modification. To test this possibility, we masked the volume of SCSα colocalized with G-3-pH and compared PGAM5 signal colocalization with SCSα colocalized with G-3-pH (lilac) to PGAM5 colocalization with SCSα signals unassociated with G-3-pH (orange) (Fig 7G). These comparisons revealed that a much smaller percentage (mean=5.7, p=0.0156) of the PGAM5 signal volume colocalized with SCSα G-3-pH as compared to the percentage of the PGAM5 signal volume colocalized with SCSα lacking association with G-3-pH (mean=18.7) (Fig 7H). Likewise, SCSα G-3-pH demonstrated less colocalization with PGAM5 as compared to SCSα lacking G-3-pH (mean= 28.7 vs. 34.9, p=0.0469, Fig 7I).

### PGAM5 phosphatase displays decreased colocalization with mitochondrial Tom20 in tumors as compared to stroma

The enrichment of pHis signals observed in stroma as compared to tumor could not be attributed to a significant elevation of ACLY G-3-pH or SCSα G-3-pH in the stroma (Fig 5) nor could it be justified by the preferential expression of a phosphatase in the stroma (Fig 6). Altered localization of a pHis phosphatase within tumor cells is one possible explanation for decreased pHis signals in tumor cells as compared to stromal cells.

Since PGAM5 can be released from damaged mitochondria through proteolytic cleavage(Sekine et al., 2012) and can dephosphorylate NME2 in the cytosol(Panda et al., 2016), we wondered whether PGAM5’s mitochondrial localization is altered in tumor (CK19+) cells as compared to stromal cells (CK19-). We triple-stained FFPE mouse KP^f/f^C PDAC advanced stage tumor sections with IHC-IF-TSA, examining colocalization of PGAM5 signals (Cy5) with signals for the outer mitochondrial membrane protein Tom20 (Cy3). CK19’s signals (fluorescein) were used to mask the tumor’s area. Super-resolution, z-series images were analyzed and revealed greater colocalization of PGAM5 with Tom20 in CK19-cells (mean =54.7) as compared to CK19+ cells (mean=47.4) (n=8, p=0.007) (Fig 8A-B). These results suggest that in response to mitochondrial damage in tumor cells, proteolytic cleavage of PGAM5 results in a portion of PGAM5 localizing to the cytoplasm, where it would have access to cytoplasmic pHis substrates.

**Fig. 8:**
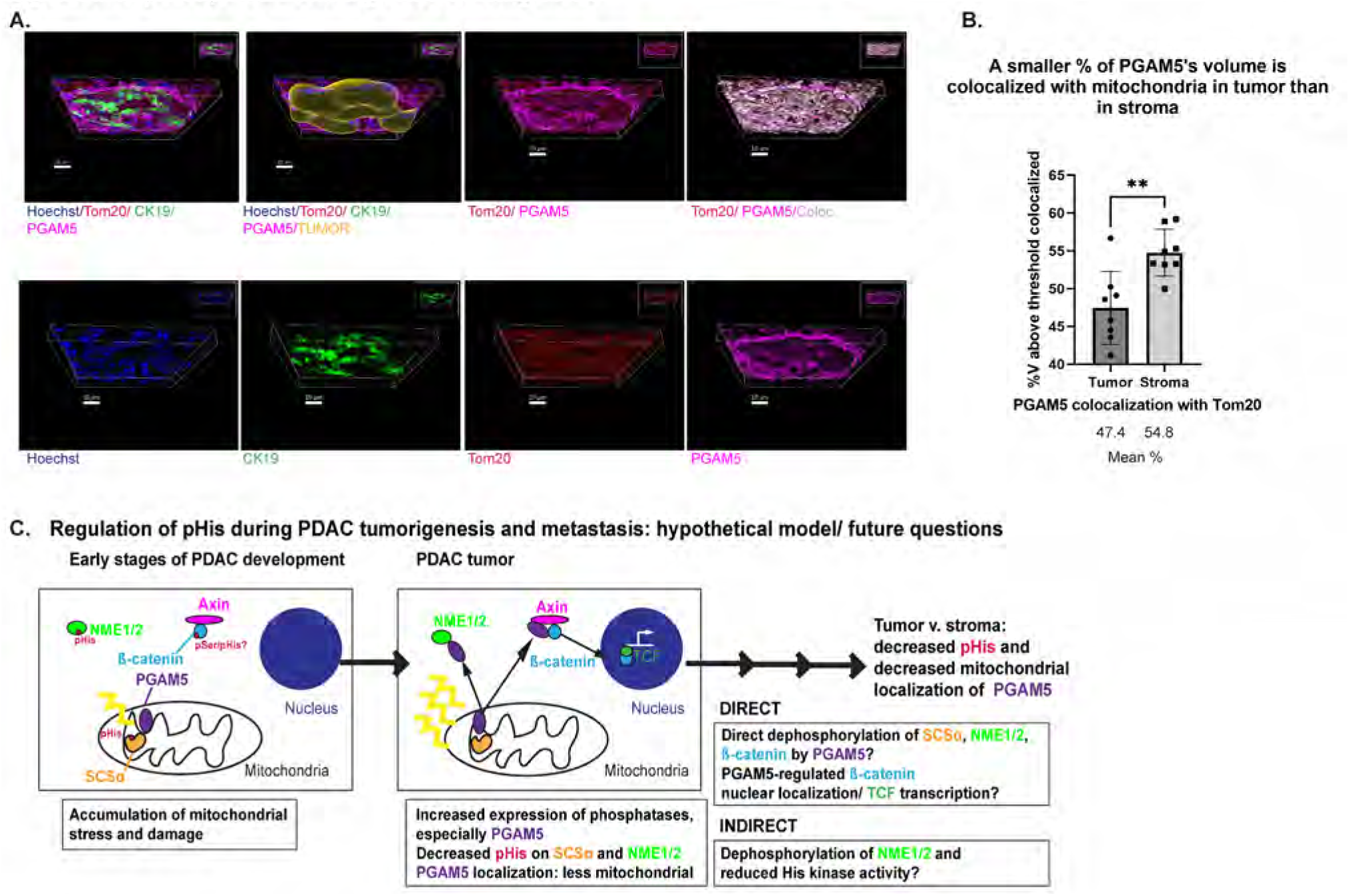
Altered mitochondrial localization of PGAM5 in tumor v. stroma suggests a hypothetical model for regulation of pHis by PGAM5 during PDAC tumorigenesis. (**a**), IHC-IF-pHis-TSA protocol (Fig S1) applied to FFPE KP^f/f^C mouse PDAC tumor serial sections analyzed with triple-staining using TSA Cy5 (PGAM5, pink signals), TSA Cy3 (Tom20, mitochondrial marker, red signals), and TSA fluorescein (CK19, green signals). Super-resolution (Airyscan, Zeiss) z-series images rendered in 3D using Imaris. Merged and colocalization channels (top); individual channel’s signals (bottom). Colocalization shown in black to white (low to high) to represent the intensity of colocalization. Scale bars are shown at the bottom left of images. (**b**) % PGAM5 volume colocalized with Tom20 in tumor or stroma (n=8, **p=0.007, Mann-Whitney). V=volume. (**c**) **Proposed model**. *Normal pancreas*: intact mitochondria, pHis=pHis levels are higher (NME1/2, SCSα) than in tumor, PGAM5 localizes to the inner and/or outer membrane of intact mitochondria. *PDAC tumorigenesis*: Increased ROS, mitochondrial fission and fragmentation are associated with PDAC tumorigenesis (Bai et al., 2020; Carmona-Carmona et al., 2022; Courtois et al., 2021; Yu et al., 2019). Our data indicate that PGAM5 expression is increased (Fig 1) and less localized with mitochondria (**a-b**) in tumor cells, and 1- and G-3-pHis (SCSα) signals are reduced in tumor tissue lysates (Fig 1) and in IHC-IF-pHis-TSA analysis (Fig 3). **Hypothesis**: Relocalization of PGAM5 due to mitochondrial damage could facilitate *DIRECT* dephosphorylation of SCSα in the mitochondrial matrix and/or known substrates of PGAM5, NME His kinase or β-catenin (pSer and/or pHis, activating its nuclear localization and TCF-mediated transcription). Alternatively, reduction in G-3-pHis signals in tumors could arise in an *INDIRECT* manner due to PGAM5’s dephosphorylation and inactivation of NME His kinase function. pHis=phosphohistidine, pSer=phosphoserine.

A model for regulation of pHis by PGAM5 during PDAC tumorigenesis based upon our data is presented in Fig 8C). The details of this model and the questions that it poses are addressed in the Discussion.

### PGAM5 is more highly expressed in human PDAC metastases as compared to primary tumors

To elucidate possible pHis targets for therapeutic modulation in human pancreatic ductal adenocarcinoma (PDAC) invasion, we compared expression of pHis and pHis regulators (Table 1) in lysates of human clinical samples, comprising 9 matched primary tumors and metastases in an immunoblotting analysis similar to Fig 1.

PGAM5 demonstrated increased expression in 6/9 samples (Fig S6D-E, p=0.05), whereas PHPT1 protein was significantly decreased in metastases in 5/9 samples (Fig S6C and S6E, p=0.03). LHPP expression was not significantly changed (average expression =1.05 in metastases relative to tumor, Fig S6F), and only one sample recorded a 2-fold change (decrease) in metastases (Table 4). However, within tumor/metastasis pairs, pHis phosphatase signals did not clearly correlate with the detected increase, on average, of pHis signals in metastases.

**Table 4.** Protein expression in metastases relative to paired human PDAC tumors (UNMC)

|  | 11 | 29 | 80 | 89 | 103 | 105 | 107 | 112 | 116 |
| --- | --- | --- | --- | --- | --- | --- | --- | --- | --- |
| **ACLY\*** | 1.08 | 0.57 | 0.63 | 0.41 | 0.32 | 0.72 | 1.38 | 0.87 | 0.94 |
| **SCSα 3-pHis\*** | 4.55 | 2.29 | 0.76 | 8.85 | 0.01 | 2.31 | 2.41 | 19.06 | 0.82 |
| **SCSα\*\*** | 2.58 | 2.65 | 4.3 | 3.32 | 0.58 | 2.28 | 0.5 | 8.22 | 0.66 |
| **PGAM5\*\*** | 2.12 | 5.92 | 18.28 | 5.17 | 10.52 | 0.95 | 0.4 | 60.16 | 0.95 |
| **PHPT1\*** | 0.88 | 0.2 | 0.52 | 0.3 | 0.49 | 0.95 | 0.74 | 0.15 | 1.42 |
| **LHPP\*** | 1.28 | 0.56 | 1.42 | 0.88 | 0.55 | 0.49 | 1.42 | 1.68 | 1.14 |
| **NME1/2 1-pHis: 17kDa\*** | 1.72 | 1.53 | 1.01 | 2.22 | 0.34 | 0.62 | 5.91 | 2.06 | 0.7 |
| **NME1/2: 17kDa\*** | 0.56 | 0.76 | 0.84 | 0.86 | 1.41 | 0.33 | 0.11 | 0.54 | 2.43 |
| **NME1/2 1-pHis: 19kDa\*** | 1.70 | 1.03 | 2.50 | 0.89 | 0.24 | 0.37 | 2.38 | 1.79 | 0.45 |
| **NME1/2: 19kDa\*** | 0.94 | 0.73 | 0.87 | 0.61 | 0.68 | 0.53 | 0.93 | 1.34 | 0.78 |

Quantification of background-subtracted signals on immunoblots normalized to GAPDH\* or α-tubulin\*\*, expression in metastases relative to tumor. ≥2-fold increase; ≤2-fold decrease. Signals with a significant increase or decrease in metastases are indicated.

Interestingly, signals for both isoforms of pHis increased in metastases as compared to tumors (Table 4), contrasting with the observed pHis expression in mouse pancreatic tumors as compared to normal pancreas (Table 2, Fig 1). For pHis signals, SCSα G-3-pH yielded the most striking result, displaying 2-fold or greater increased expression in 6/9 metastases, and a similar increase in levels of total SCSα protein was observed (Fig S6A, 4.6-fold and S6E, 2.8-fold, p=0.05). Although anti-1-pHis signal differences were not statistically significant, these signals were elevated greater than 2-fold in 4/9 metastases for either a 17 kDa or 19 kDa band expected to be NME1/2, and these changes did not correspond to altered total expression of NME1/2 proteins (Fig S6B and S6F). ACLY expression was decreased in metastases (average expression =0.77 in metastases relative to tumor, Fig S6F). ACLY G-3-pH expression was obscured by background on blots and was not quantitated.

## Discussion

Regulators and substrates of His phosphorylation have been previously linked to roles in tumorigenesis (ACLY(Carrer et al., 2019; Shah et al., 2016), LHPP(Hou et al., 2020; Li et al., 2019; Sun et al., 2020; Wang et al., 2021; F. Wu et al., 2020; Zhang et al., 2020; Zheng et al., 2018), PGAM5(Cheng et al., 2018; Ng Kee Kwong et al., 2018)), and metastasis (ACLY(Wen et al., 2019), PHPT1(Xu et al., 2010), PGAM5(Kang et al., 2015), NME1(Kantor et al., 1993; Khan et al., 2019; Lee et al., 2018; Steeg et al., 1988; Suzuki et al., 2004)) and localize to varying subcellular regions, based upon data from this study and previous publications (detailed in Table 1). Analyzing these proteins and pHis signals in mouse pancreatic tumors, we found that pHis signals were decreased as compared to normal pancreatic tissue and non-tumor or stromal cells. Immunoblots comparing lysates of mouse pancreatic tumors to normal pancreas revealed a decrease in pHis and total pHis protein expression (Fig 1, NME1/2, SCSα). Using the IHC-IF-pHis TSA protocol that we developed to analyze mouse FFPE PDAC tumors, we demonstrated an enrichment of both 1-pHis (likely NME1/2) and G-3-pHis signals in stroma (CK19-) rather than tumor cells (Fig 3, Fig S2).

Increased expression of pHis phosphatases in mouse pancreatic tumors could in principle account for the observed reduction in pHis levels. Decreased pHis levels in mouse PDAC tumor lysates as compared to normal pancreas lysates on immunoblots correlated with increased pHis phosphatase expression (PGAM5, PHPT1, LHPP) (Fig 1). Colocalization between G-3-pH signals and pHis enzymes ACLY and SCSα was reduced on average in tumor areas as compared to stromal areas (Fig 5). Signals for PHPT1 and PGAM5 exhibited inverse correlation coefficients when colocalized with G-3-pH signals in 3D analyses of mouse PDAC tumor FFPE sections. However, neither pHis phosphatase was enriched in tumor regions as compared to stroma (Fig 6). PGAM5’s localization to mitochondria was reduced in tumor cells as compared to stroma cells, suggesting that cytoplasmic forms might dephosphorylate extramitochondrial pHis substrates in tumor cells as compared to stroma cells, leading to enrichment of pHis signals in the stroma (Fig 8).

We sought to identify the source of the enriched pHis signals in the stroma. We focused our detailed analyses on proteins possessing Gly-3-pHis motifs (ACLY, SCSα) by using the sequence and isoform-specific anti-3-pHis mAb, SC44-1, since these enzymes use a 3-pHis enzyme intermediate and are well-expressed in PDAC. We compared G-3-pHis signal expression in FFPE tissue processed using the IHC-IF-pHis-TSA protocol (Fig 3) to anti-G-3-pHis immunoblots of tumor, immune, and CAFs populations isolated by FACS (Fig 4). Although the most abundant G-3-pHis signals were found in non-tumor cells for both analyses, G-3-pHis signals for CAFs were better detected using IHC-IF-TSA protocol as compared to FACS/immunoblotting. This likely reflects differences in the tumor preparation (tissue dissociation vs. FFPE) and the different types of CAFs that were analyzed by the two techniques (podplanin+ vs. αSMA+).

FACS/immunoblot analyses revealed the presence of novel forms of G-3-pHis+ ACLY protein in TAMs (Fig 4). Although the role of ACLY in myeloid cells has been explored in the literature (Covarrubias et al., 2016; de Goede et al., 2021; Lauterbach et al., 2019; Namgaladze et al., 2018; Rhee et al., 2019; Verberk et al., 2021), the expression of non-canonical ACLY isoforms in CD11b+Ly6G-myeloid cells has not been previously described and the origin and function of these isoforms is the subject of ongoing research.

We also identified an enrichment of cell-cell adhesion proteins in the putative G-3-pHis proteome of TAMs from mouse pancreatic tumors (Fig 4). Except for plakophilin-1 (Pkp1, 80.9 kDa), these proteins were identified in both tumor and TAM lysates. For example, this study is the first to predict G-3-pHis on β-catenin, and we were able to detect a small percentage of this protein in immunoblots of immunoprecipitates made with a G-3-pH-specific mAb in both sorted tumor cells from KP^f/f^C tumors and in cultured cells established from a KP^f/+^C tumor.

Interestingly, within its C-terminal transactivation domain, β-catenin possesses a GHAQ motif, which is similar to the GHAG motif of ACLY and SCSα. Identification of pHis sites and site-directed mutagenesis and are needed to validate His phosphorylation of this and other GH motifs in cell-cell adhesion proteins, and further studies with pHis site mutants could reveal a role for β-catenin and cell-cell adhesion in pHis signaling pathways. Nevertheless, based upon the quantities of peptides observed in our IP/MS studies (e.g. nearly 50-fold more peptides for ACLY than β-catenin, Supplemental File 1), we expect that overall level of histidine phosphorylation of cell-cell adhesion proteins is significantly less than that of enzymes such as ACLY, SCSα and NME1/2. This may explain why direct MS approaches to mapping pHis sites have failed to identify and validate pHis on signaling proteins due to their relative paucity in lysates as compared to pHis enzymes and the acidic conditions used in this process (Cui et al., 2021; Gao et al., 2019; Hardman et al., 2019; Hu et al., 2020; Keven Adam, 2019; Leijten et al., 2022; Zhao et al., 2021).

Our results suggest that PGAM5 is less mitochondrially localized in pancreatic tumor cells due to mitochondrial stress (Fig 6). PGAM5 localization and function is linked to the mitochondrial stress response in the literature (Bernkopf et al., 2018; Lu et al., 2014; Wang et al., 2012; Yu et al., 2020). PGAM5’s normal localization has been a topic of debate (Cheng et al., 2021), but evidence supports its localization at the inner (Lu et al., 2014; Sekine et al., 2012) and outer (Yamaguchi et al., 2019) mitochondrial membrane, or both inner and outer membranes (at contact sites or through shuttling) (Sugo et al., 2018). PGAM5 is expressed as both short (S) and long (L) isoforms(Lo & Hannink, 2006). Both are required for necroptosis(Wang et al., 2012) and can be released from damaged mitochondria by proteolytic cleavage, leading to their cytosolic localization(Sekine et al., 2012). PGAM5 dephosphorylation of pSer in dynamin-related protein 1 (Drp1) activates its GTPase and mitochondrial fission during mitochondrial dysfunction(Park et al., 2018; Wang et al., 2012). Dephosphorylation of the FUNDC1 receptor by PGAM5 regulates autophagy(Chen et al., 2014; Wu et al., 2014). PGAM5 also affects mitophagy by recruiting and stabilizing PINK1 at damaged mitochondria(Lu et al., 2014). PGAM5’s cleaved form released from damaged mitochondria increases mitochondrial numbers and dephosphorylates pSer on β-catenin, stabilizing its expression(Bernkopf et al., 2018). Additional studies would be required to map the relevant PGAM5 isoforms expressed in PDAC and validate which forms are cleaved and whether this changes during tumorigenesis and metastasis, and also determine which pHis proteins can serve as direct substrates for PGAM5.

We present a model for the regulation of pHis by PGAM5 during pancreatic tumorigenesis (Fig 8C). Evidence supports a correlation between mitochondrial fission and fragmentation with tumorigenesis and invasiveness in pancreatic cancer(Bai et al., 2020; Carmona-Carmona et al., 2022; Courtois et al., 2021; Yu et al., 2019). We propose that in normal pancreas, NME, SCSα, and β-catenin contain basal levels of pHis. Mitochondrial damage in pancreatic tumor cells leads to PGAM5’s proteolytic cleavage and some fraction of cleaved PGAM5 localizes to the cytoplasm. Here, PGAM5 can access two of its known substrates: NME2(Panda et al., 2016) and β-catenin (its known pSer substrate and a possible pHis substrate(Bernkopf et al., 2018)). There is no direct biochemical evidence that PGAM5 can dephosphorylate pHis SCSα, but our finding that PGAM5 exhibited greater colocalization with SCSα lacking a G-3-pHis signal (Fig 7), suggests that PGAM5 somehow influences the level of SCSα His phosphorylation. Whether this is a direct or indirect effect, e.g. resulting from decreased pHis NME1/2 levels, remains to be determined by in vitro biochemical analyses.

PGAM5 could contribute either directly (by dephosphorylation of NME and G-3-pHis substrates) or indirectly (by dephosphorylation and inactivation of NME His kinase activity) to the phenotypes of decreased pHis correlated with mouse PDAC tumors that we observed in this study. Although PGAM5 is histidine phosphorylated through its phospho-acceptor catalytic histidine site (Panda et al., 2016), the anti-pHis antibodies used in this study failed to detect this modification.

Decreased NME expression and His kinase functions could contribute to PDAC tumorigenesis in a PGAM5-dependent or -independent manner. Possible NME1 substrates in neuroblastoma, a background in which NME1 is over-expressed, have been identified(Adam, Lesperance, et al., 2020) and are currently being characterized in the lab for their impact upon tumorigenesis. Colocalization analyses of PGAM5, NME1, NME2, and His kinase substrates could elucidate differences in pHis wiring in PDAC vs. neuroblastoma.

Future metabolomic studies would be useful in linking histidine phosphorylation state to metabolite production. However, we expect that the decreased levels of pHis on NME and SCSα in PDAC tumor tissue lysates as compared to normal pancreas are associated with their decreased enzymatic function. Total protein expression for SCSα and NME1 was similarly decreased in mouse PDAC tumors. This implies decreased reliance upon the TCA cycle (with respect to decreased SCSα 3-pHis) and NDPK/His kinase function (with respect to decreased levels of NME 1-pHis). PGAM5 dephosphorylation of pHis substrates could mediate their degradation through ubiquitin ligase recruitment, as in PGAM5’s regulation of Bcl-XL(Niture & Jaiswal, 2011). Nevertheless, additional studies are needed to clarify the nature of these relationships.

PGAM5 presents a possible target in pancreatic tumorigenesis and metastasis. In comparisons of human PDAC metastases to matched primary tumors (Fig S6), PGAM5 expression was increased, concordant with its increased expression in mouse PDAC tumors as compared to normal pancreas (Fig 1). Although PGAM5’s contribution to pancreatic tumorigenesis has not been directly tested, high levels of PGAM5 are associated with a poor prognosis in hepatocellular carcinoma, and PGAM5 loss is associated with chemosensitivity in vitro and in vivo in a mouse model(Cheng et al., 2018). A novel PGAM5 inhibitor (LFPH-1c) was recently described but has not been tested in cancer models(Gao et al., 2021).

Additional studies are needed to elucidate the regulation of pHis in the PDAC metastasis microenvironment. Expression of SCSα (pHis and total protein) was increased in human PDAC metastases as compared to primary tumors, contrasting with its reduction in comparisons of mouse PDAC tumors to normal pancreas. SCSα G-3-pHis localizes to the mitochondrial matrix and participates in the TCA cycle utilizing a G-3-pHis intermediate to synthesize succinate and ATP from succinyl CoA(Huang & Fraser, 2021). Recent data suggests that succinate accumulation alters cellular epigenetic states to promote tumorigenesis and metastasis(Aspuria et al., 2014; Letouze et al., 2013; Wang et al., 2016; J. Y. Wu et al., 2020; Zhao et al., 2017).

Moreover, SCSα G-3-pHis-mediated synthesis of ATP through mitochondrial substrate-level phosphorylation in this reaction could compensate for reduced oxidative phosphorylation in tumor cells(Seyfried et al., 2020). Whether increased SCSα G-3-pHis expression in PDAC metastases correlates with increased activity and succinate or ATP synthesis poses an interesting question. Proximity ligation studies of PGAM5 and SCSα G-3-pH in metastases as compared to tumors could reveal whether or not PGAM5’s distance from SCSα precludes its regulation of pHis on SCSα in this environment. Biochemical analyses of PGAM5’s impact upon SCS enzymatic functions, and metabolomic analyses of succinate levels in PDAC tumors and metastases could also provide insight into these processes.

Overall, we describe a technique to broadly analyze pHis protein localization in the context of substrates and His kinase/pHis phosphatase regulators in PDAC tumor tissue and other tissue models for human disease. By circumventing acid- and heat-sensitivity of pHis, the IHC-IF-pHis-TSA protocol provides a starting point for future spatial proteomics studies that have the potential to localize pHis signals, regulators, and substrates to specific cell populations to better understand how these networks are regulated and discover how regulation changes with respect to cancer and other pathologies. Future studies are needed to determine whether clinical specimens can be analyzed in a similar manner. Ultimately, these studies could identify additional pHis regulation targets for therapeutic development or pHis regulation signatures for early detection of tumor development.

## Materials and Methods

**Table S1.**
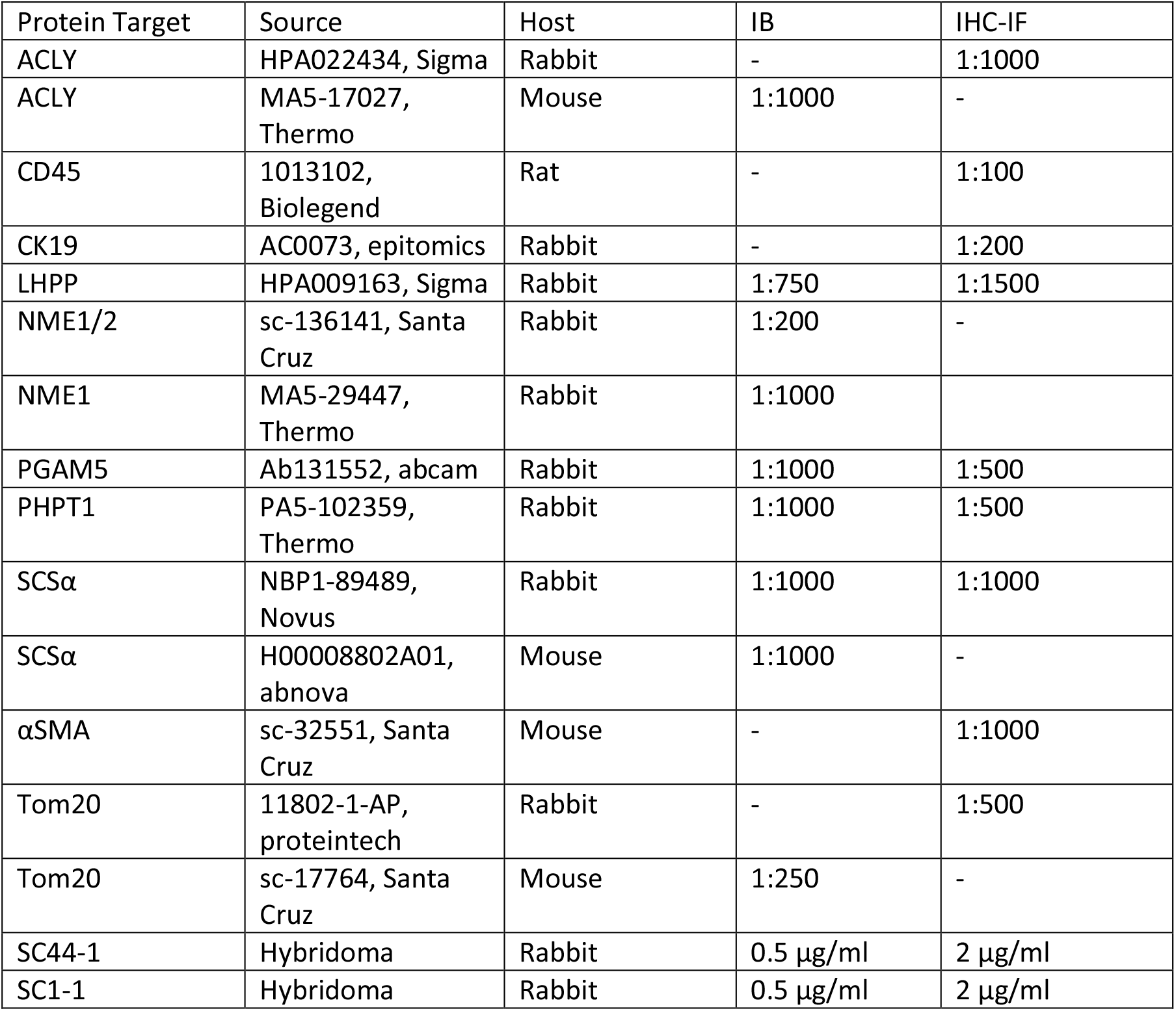
Antibodies, Table S1

### Mice

Except where indicated, PDAC tumors analyzed were harvested from KP^f/f^C mice (Kras^LSL-G12D/+^; Trp53^flox/flox^; Pdx1-Cre mice on FVB background, characterized in a previous publication(Shi et al., 2019)). Normal pancreas littermate controls (P^f/f^C, K^LSL-G12D/+^) were isolated for comparisons. KP^f/+^C (Kras^LSL-G12D/+^; Trp53^flox/+^; Ptf1a-Cre on C57BL/6) and KC (Kras^LSL-G12D/+^; Ptf1a-Cre on C57BL/6) PDAC tumors were kindly provided by G. Wahl (Salk) and included to demonstrate the existence of certain PDAC phenotypes in multiple backgrounds. The orthotopic pancreatic tumor shown (Fig 4C) was generated from cells established from KP^f/+^C mice. No sexual dimorphism was noted in any mouse model, and therefore both males and females are represented in all data sets. Mice were bred and maintained in the animal care facilities at the Salk Institute. Genotypes of individual mice were determined by PCR using genomic DNA from tail biopsies with Bioline MyTaqTM Extract-PCR Kit. Standard PCR per manufacturer’s protocol was performed, annealing at 58 or 60 °C for 35 cycles. Primer sequences are available upon request. All animal experiments were performed according to protocols approved by the Salk Institute Animal Care and Use Committee.

### Cell culture

1245 cells were established from a KP^f/+^C (Kras^LSL-G12D/+^; Trp53^flox/+^; Ptf1a-Cre on C57BL/6) and are suitable for orthotopic mouse experiments. For preparation of lysates used in the IP of β-catenin, we cultured these cells at 37°C 5% CO2 in a humidified chamber on tissue-culture treated polystyrene plates using DMEM+10% FBS with antibiotics added as media.

### Human specimens

De-identified, paired samples (9) for primary pancreatic tumors and metastases, prepared upon autopsy from patients who had given prior informed consent, were provided as frozen tissue by M.A. Hollingsworth (UNMC) with approval by Salk Institutional Review Board (IRB), protocol #17-0005. Frozen tissue specimens were homogenized in cold Tissue Extraction Reagent II (Invitrogen) containing protease and phosphatase inhibitors (Complete mini and PhosSTOP, Roche) at 100 mg tissue per 1 mL and sonicated as previously(Shi et al., 2019). Equal amounts of proteins were resolved by SDS-PAGE, transferred to PVDF membranes and immunoblotted.

### Tissue dissociation and FACS

Tumors were rapidly isolated from CO_2_-euthanized mice and dissociated by incubating in 0.2 mg/ml pronase (Roche), 0.01 mg/ml DNase I, and 1 mg/ml collagenase P (Sigma) at 37°C for 50 min with shaking and pipetting. Suspensions were washed with FACS buffer (PBS +0.5% BSA+ 0.5mM EDTA) and filtered (100 µm). Pelleted cells were lysed in ACK lysis (Thermo) to exclude red blood cells from the analysis and washed in FACS buffer. Single cell suspensions were incubated on ice with mouse Fc receptor block (BD Biosciences, 1:200) followed by antigen-specific antibodies in FACS buffer. PI (molecular probes, 1:1000) was used to exclude dead and dying cells. Cells were labeled with CD45 (BV421), EpCAM (Alexa Fluor 647), CD11c (PE-Cy7), CD11b (PE), Ly6G (APC-Cy7) and CD3 (BV785) (Biolegend, 1:200) antibodies. Fluorescence minus one (FMO) staining controls were included for gating populations of interest. Cells were FACS purified at the Salk Institute’s Flow Cytometry core facility on a BD Biosciences Aria cell sorter (100-μm size nozzle, 1 × PBS sheath buffer with sheath pressure set to 20 PSI). Cells were sorted into 0.5 ml of FACS buffer and maintained on ice until pelleting at 500xg for 5 min at 4°C. Pellets were suspended in lysis buffer (10 mM Tris-HCl pHis 8.8; 0.1% SDS; 1% sodium deoxycholate; 0.5 mM EDTA pHis 8; 150 mM NaCl with 1 mM PMSF and complete mini EDTA-free protease inhibitor cocktail and PhosSTOP added) using volumes to produce an equal number of cells per ml for each population and stored at −80°C. Lysates were cleared at 20,000xg for 10 min upon thawing on ice and equal volumes were processed using SDS-PAGE, transfer, and immunoblotting.

### Tissue processing and embedding

The fixed tissues were processed on a Sakura Tissue Tek VIP ™ tissue processor in a shortened cycle. The steps are as follows:

1. Two changes of 70% reagent alcohol (Fisher cat # HC1500), for a total of 30 min at room temperature with gentle stirring.
2. Two changes of 95% reagent alcohol (Fisher cat # HC1300) for a total of 25 min at room temperature with gentle stirring.
3. Three changes of 100% reagent alcohol (Fisher cat # HCHC600) for a total of 30 min at room temperature with gentle stirring.
4. Three changes of Clear-Rite 3TM (xylene substitute, Epredia cat # 6915) for a total of 20 min at room temperature with gentle stirring.
5. Three changes of Histoplast PE paraffin (Epredia cat # 8330) at 62°C for a total of 60 min with gentle stirring.

The cassettes were then transferred to the Thermo Scientific HistostarTM embedding station and embedded with paraffin, Type L (Epredia cat # 8330) at 60°C and then cooled at 4°C to solidify.

### Histological preparation and staining

Preparation was modified from previously published protocols(DelGiorno et al., 2020; Shi et al., 2019) to conserve pHis signals for multiplexed staining. Rapidly isolated, thinly sliced (1-2 mm) tissue samples were fixed overnight at 4°C in neutral-buffered formalin (Fisher Scientific) with PhosSTOP added (Roche), embedded in paraffin using a shorter cycle (see ‘Tissue processing and embedding’). These fragile FFPE blocks were incubated on ice prior to sectioning, and 5 µm sections were placed in a 36C water bath and mounted to slides. After drying slides overnight at room temperature, sections were deparaffinized in xylene, rehydrated in a series of ethanol (2×100%, 2×95%, 75%, 50%) and then washed twice in MilliQ H_2_O. Endogenous peroxidase activity was blocked with a 3% H2O2 for 10 min. After a MilliQ H_2_O wash, tissue was outlined using a hydrophobic pen (ImmEdge, Vector Labs), then incubated in TBS + 0.05% Tween 20 (TBST) for 5 min. Slides stained with antibodies other than anti-pHis were treated with microwave AR (10 mM sodium citrate, 0.05% Tween20 pHis 6.0) treatment (10 min in boiling buffer), followed by washes in MilliQ H_2_O and TBST. Sections were blocked with blocking buffer (TBST + 5% bovine serum albumin) for 45 min at room temperature. Primary antibodies were diluted in blocking buffer and incubated for 2 hr at room temperature (anti-pHis) or overnight at 4°C (other antibodies). Information on primary antibodies is provided in Supplementary Table S1. Slides were then washed (TBST) and incubated in HRP based secondary antibodies, either Signal Stain Boost (for primary antibodies of mouse or rabbit origin, CST) or ImmPRESS® HRP Goat Anti-Rat IgG, Mouse adsorbed Polymer Detection Kit, Peroxidase, (Vector). IHC-IF-pHis-TSA staining utilized TSA treatment (1:50) for 8 min using either Cy5, Cy3, or fluorescein (Akoya Biosciences), following TBST washes after secondary antibody staining. Subsequently, TBST washes were applied. For multiplexing of signals, an additional AR treatment, block, and overnight incubation in primary antibody were followed. Secondary antibodies and TSA staining steps were performed on the following day, and this process was repeated for a third day to obtain triple-staining (Cy5, Cy3, fluorescein). Once all fluorophore staining was completed, slides were stained for nuclei using Hoechst 33342 in PBS (2 µg/ml incubated for 20 min at room temperature), followed by PBS washes and mounting with Prolong Gold Antifade (Thermo).

Chromogenic staining (Fig S1) followed a well-detailed, published protocol(Luhtala & Hunter, 2020). Hematoxylin and eosin (H&E) staining was performed to assess tissue morphology for chromogenic staining.

### Immunofluorescent (IF) and chromogenic IHC images: capture and analysis

All slides were scanned and imaged (single plane) on an Olympus VS-120 Virtual Slide Scanning microscope (brightfield and fluorescence). 2D images were analyzed using QuPath. Tissue sections were outlined (polygon tool) to create annotations, cells were detected using DAPI signal with default settings. Classifications (e.g., CK19+, αSMA+) were defined from channels, and classifiers were defined by single threshold for mean cell signals, selected to exclude background signals in controls. Classifiers were combined to generate percentages of single and double-positive cells within samples.

For super-resolution 3D analyses, images of z-sections were captured using the Zeiss LSM 880 Airyscan with optimal frame size and optimal z intervals (Piezo z-stage) selected; resultant images were deconvolved by Airyscan processing (Zeiss). Imaris software (v9.8.0, Oxford Instruments) was used for all 3D analyses and rendered images. Control images were captured, and thresholds for colocalization and min and max signals used for each channel were selected to eliminate control signals. Channels were processed on Imaris using a 1 µm filter for background subtraction of Cy3, Cy5, and fluorescein signals; DAPI background subtraction used a 3µm filter. Masks were created for tumor (CK19+) signals with settings for threshold and smoothing selected to best encompass tumor cells within all images being compared. Voxels were set to 0 for signals outside of the masked region for “tumor” and within the masked region for “stroma”. To analyze SCSα G-3-pHis signals, a mask was created without smoothing from the SCSα / G-3-pHis colocalization channel. Voxels were set to 0 for signals outside of the masked region to analyze SCSα G-3-pHis signals, and voxels were set to 0 for signals outside of the masked region to analyze SCSα signals not associated with G-3-pHis.

### SDS-PAGE, immunoblotting (IB), and quantitation

We made slight modifications to a protocol that optimizes pHis conservation for SDS-PAGE(Kalagiri et al., 2020). 5X pHis 8.8 loading buffer (20% SDS; 50 mM Tris-HCl pH 8.8; 100 mM DTT; 10% glycerol; 10 mM EDTA pH 8; bromophenol blue) (for pHis conservation) or 2X loading buffer (4% SDS; 20% glycerol; 100 mM Tris; 40% β-mercaptoethanol) (for boiled IP samples) was used, and samples were resolved on polyacrylamide gels consisting of a separating gel [7.5%, 12.5%, or 15% (w/v) acrylamide, 0.1% (w/v) bisacrylamide, 0.38 M Tris-Cl, pH 8.8; 0.1% (w/v) SDS] and a stacking gel [4% (w/v) acrylamide; 0.1% (w/v) bisacrylamide; 0.125 M Tris-Cl, pH 8.8] for gels requiring pHis conservation and pH 6.8 for boiled IP samples, 0.1% (w/v) SDS] at constant voltage (105 V), and gels were transferred overnight at 4 °C with constant voltage (22 V) to PVDF membranes optimized for Odyssey detection. Li-Cor Odyssey protocols were followed for blotting of membranes with primary and secondary antibodies (Alexa dye 680 and IRDye 800), and after washing, these were scanned to the Li-Cor Odyssey for quantitative detection of signals. Multi-antibody staining for multiple proteins within a single IB was performed by utilizing both detection channels (680 and 800) and sequentially immunoblotting for non-overlapping signals without the use of antibody dissociation. Careful analysis of bands observed was conducted to avoid confusing background with *bona fide* signals; anti-pHis blotting was always performed first to optimize detection. For quantitation, bands for proteins of interest were manually defined using a rectangle or ellipse, and an identical size/shape background band was manually defined (user defined background method). Normalized intensities were calculated from I.I.K counts given by the Li-Cor Odyssey software by subtracting background values from bands of interest and calculating this signal as a fraction of background-subtracted signal for a normalizing protein.

### Immunoprecipitation (IP)

PDAC tumors from three male 40-day old KP^f/f^C mice were dissociated and sorted for tumor (EpCAM+CD45-) and TAMs (CD45+CD11b+Ly6G-) as detailed in ‘Tissue Dissociation and FACS’. All buffers were pre-chilled with 1 mM PMSF and complete mini EDTA-free protease inhibitor cocktail and PhosSTOP added, and all steps were performed on ice in a room at 4°C. Cells (3×10^5^ per IP per population) were lysed in IP lysis buffer (50 mM Tris-HCl pH7.5; 150 mM NaCl; 1% SDS; 0.5% sodium deoxycholate), with needle shearing of chromatin (27-g). Cells were diluted 10-fold using IP dilution buffer (50 mM Tris-HCl pH7.5; 150 mM NaCl) and incubated with rotation for one hour with antibody (SC44-1 anti-G-3-pHis, 6 µg), then washed protein A agarose beads were added. Rotation with beads continued for 5 hr, then the unbound fraction was removed, and beads were washed 3x in IP dilution buffer. An equal volume of buffer (100 µl) was left remaining for all samples, and 2X loading buffer was added in that volume. Samples were boiled for 10 min, and 20 µl was loaded for SDS-PAGE and immunoblot analysis. The remainder was submitted for mass spectrometry. This protocol was slightly modified for the IP of β-catenin from 1245 KP^f/+^C cell lysates in that 4×10^6^ cells per IP were used.

### Controls used in IHC, IHC-IF, IP, and IB

Prior to including an experiment in quantitation for IHC or IHC-IF, we compared pHis peptide or 1- or 3-pTza (Fuhs et al., 2015; Kee et al., 2010) block of antibody or antigen retrieval and compared signals for a serial section stained with antibody not pre-treated with block or antigen retrieval. Generating pHis-phosphorylated peptides using phosphoramidate and their use as a block to pHis binding of antibodies for IHC experiments has been described previously (Luhtala & Hunter, 2020). Images were scaled using the same min to max values, and an experiment was only included in our analyses if signals were eliminated in the control. Sections stained with only primary antibodies served as a control for other antibodies.

For the IP-anti-G-3-pHis controls, one control IP included SC44-1 anti-G-3-pHis antibody pre-incubated 30 min at room temperature with 5 µg of AGAG 3-pTza AGAG peptide (Fuhs et al., 2015); another control IP used rabbit IgG in place of antibody (6 µg).

For IB acid/boiling/neutralization treatment to eliminate pHis on protein in lysates, samples were boiled for 10 min in 1X pH 8.8 buffer with glacial acetic acid added to 1%, then neutralized with sodium hydroxide added to 0.4 M.

### Mass spectrometry

Samples were precipitated by methanol/ chloroform and redissolved in 8 M urea/100 mM TEAB, pH 8.5. Proteins were reduced with 5 mM tris(2-carboxyethyl) phosphine hydrochloride (TCEP, Sigma-Aldrich) and alkylated with 10 mM chloroacetamide (Sigma-Aldrich). Proteins were digested overnight at 37°C in 2 M urea/100 mM TEAB, pH 8.5, with trypsin (Promega). Digestion was quenched with formic acid, 5 % final concentration.

The digested samples were analyzed on a Orbitrap Eclipse tribrid mass spectrometer (Thermo). The digest was injected directly onto a 25 cm, 100 um ID column packed with BEH 1.7um C18 resin (Waters). Samples were separated at a flow rate of 300 nl/min on a nLC 1000 (Thermo). Buffer A and B were 0.1% formic acid in water and 0.1% formic acid in 90% acetonitrile, respectively. A gradient of 1-25% B over 100 min, an increase to 40% B over 20 min, an increase to 90% B over 10 min and held at 90%B for a final 10 min was used for 140 min total run time. Column was re-equilibrated with 20 ul of buffer A prior to the injection of sample.

Peptides were eluted directly from the tip of the column and nanosprayed directly into the mass spectrometer by application of 2.5 kV voltage at the back of the column. The Orbitrap Eclipse was operated in a data dependent mode. Full MS scans were collected in the Orbitrap at 120K resolution with a mass range of 380 to 1800 m/z and an AGC target of 1e6. The cycle time was set to 3 sec, and within this 3 sec the most abundant ions per scan were selected for HCD MS/MS (NCE 35) in the Orbitrap at 60K resolution with an AGC target of 1e5 and minimum intensity of 5000. Maximum fill times were set to 246 ms and 8000 ms for MS and MS/MS scans respectively. Quadrupole isolation at 0.7 m/z was used, monoisotopic precursor selection was enabled and dynamic exclusion was used with exclusion duration of 50 sec.

Protein and peptide identification were done with Integrated Proteomics Pipeline – IP2 (Integrated Proteomics Applications). Tandem mass spectra were extracted from raw files using RawConverter (He et al., 2015) and searched with ProLuCID (Xu et al., 2015) against Uniprot mouse database. The search space included all fully-tryptic and half-tryptic peptide candidates. Carbamidomethylation on cysteine was considered as a static modification. Data was searched with 50 ppm precursor ion tolerance and 600 ppm fragment ion tolerance. Identified proteins were filtered to using DTASelect (Tabb et al., 2002) and utilizing a target-decoy database search strategy to control the false discovery rate to 1% at the protein level (Peng et al., 2003).

### Statistical and functional analyses

All graphs and statistical comparisons included were performed using Prism (GraphPad, v9). Non-parametric two-tailed testing at the p≥0.05 level of significance was tested with either Mann-Whitney or Wilcoxon signed rank for unpaired and paired data, respectively. Individual values are graphed as symbols with standard error bars and mean represented.

To functionally annotate the putative G-3-pHis proteome of TAMs, we filtered results for peptides obtained from IP-MS analysis as described (Supplementary File 1, ‘Legend’). Proteins predicted in greater abundance based on the number of peptides in IP-anti-G-3-pHis of TAMs, but not controls, were scanned for GH motifs. A list of these proteins was submitted for functional annotation using The Database for Annotation, Visualization, and Integrated Discovery (DAVID) v2021 (Huang da et al., 2009a, 2009b; Sherman et al., 2022), and a gene ontology biological process (GO_BP_FAT) report was generated for this group of proteins (Supplemental File 1, ‘GO_BP_FAT-GH-TAMs’).

## Acknowledgements

We thank Dr. Shixin Ma for valuable discussions related to ACLY expression in immune cells. We thank Lynsey Hamilton (Imaris) and Drs. Uri Manor, Rebecca Gilson, and Cayla Miller in the Waitt Advanced Biophotonics Core at Salk for providing the facilities and training for slide scanning, super-resolution microscopy, and Imaris analysis. We acknowledge Drs. James Moresco, Jolene Diedrich and Antonio Pinto for their contributions to the design, execution, and analyses of mass spectrometry experiments (Salk Mass Spectrometry Core). We are grateful to Dr. Kimberly McIntyre and the UCSD Tissue Technology Shared Resource staff for aiding in the development and execution of the FFPE tissue embedding protocol. We also thank Dr. Walter Eckhart and all members of the Hunter laboratory for feedback on this project and the development of this manuscript.

## Funding

This study was supported by NIH grants CA080100, CA082683, CA194584 and CA242443 to T.H.; CA197687 to G.M.W.; P50CA127297, U01CA210240, P30CA36727 and 5R50CA211462 to M.A.H.; a Lustgarten Foundation for Pancreatic Cancer Research Dedicated Program Grant to T.H; and Freeberg Foundation funding to G.M.W. T.H. is the Renato Dulbecco Chair in Cancer Research and a Frank and Else Schilling American Cancer Society Professor; G.M.W. is the Daniel and Martina Lewis Chair. N.E.L. was supported by a George E. Hewitt Foundation Postdoctoral Fellowship and an American Cancer Society Postdoctoral Fellowship 127795-PF-15-030-01-CSM, and N.K.L. was supported by the Salk Institute Training Grant T32 CA009370, a Hope Funds for Cancer Research Postdoctoral Fellowship (HFCR-20-03-03), and a Sky Foundation Seed Grant. K.E.D. is supported by a Vanderbilt-Ingram Cancer Center Support Grant Development Award (5P30 CA068485-26), a Vanderbilt Digestive Disease Research Center Pilot and Feasiblity Award (5P30 DK058404-20), an American Gastroenterological Association Research Scholar Award (AGA2021–13–02), NIH-NIGMS 1R35 GM142709–02, and a Sky Foundation Seed Grant (AWD00000079). S.M.K. is supported by NIH R01CA240909 to S.M.K. and G.M.W., with previous support by NIH R21AI151986. The Salk Institute Waitt Advanced Biophotonics Core, the Flow Cytometry Core and the Mass Spectrometry Core are supported by a National Cancer Institute Cancer Center Support Grant (CCSG P30CA014195) and the Helmsley Center for Genomic Medicine. The UCSD Tissue Technology Shared Resource is supported by a National Cancer Institute Cancer Center Support Grant (CCSG P30CA023100). The funders had no role in the study design, data collection and analysis, decision to publish, or preparation of the manuscript.

## Author Contributions

All authors participated in the final approval of the manuscript. N.E.L. conceived and designed the experiments, performed data acquisition, analysis and interpretation, drafted the article, and participated in the revision of the article for publication. T.H. contributed to the design of the experiments, interpretation of data, and revision of the article for publication. N.K.L. designed, executed, and analyzed FACS experiments, provided a supplemental figure and contributed to the revision of the manuscript. K.E.D. guided the development of IHC protocols and participated in the revision of the manuscript. Y.S. provided KP^f/f^C mice and tissue lysates for human primary PDAC tumors and metastases, from samples obtained from M.A.H. R.N. executed tissue sectioning and mounting of FFPE blocks. S.M.K. provided valuable advice in the development of immune cell experiments and provided antibodies for FACS labeling. G.M.W. provided important feedback in the development of the project, lent personnel to support the project, and provided the microtome used in sectioning.

## Competing Interests

The authors have no competing interests to declare.

## Supplemental File Legend

**Supplementary File 01**. *IP-SC44-1 anti-G-3-pHis-MS results*. **Legend**. Description of how results are filtered and represented in the sheets provided. **Unfiltered Results**. Results are provided without filtering; proteins selected for analysis are in boldface with font color described in the legend. **Compare-to-prior-results**. Predicted GpH proteins were compared to prior analyses (Adam, Lesperance, et al., 2020; Fuhs et al., 2015; Hindupur et al., 2018) using Bioinformatics and Research Computing Tools (http://barc.wi.mit.edu/tools/compare/). **Tumor-only-motifs**. Peptides predicted only in IP from tumor lysates that contained GH motifs are listed along with their size and location of GH motifs in mouse and human proteins. **Tumor-only-pept-count**. Peptide counts for tumor only proteins in all IP samples are provided. **TAMs only-motifs**. Peptides predicted only in IP from tumor-associated monocytes/macrophages (TAMs) lysates that contained GH motifs are listed along with their size and location of GH motifs in mouse and human proteins. **TAMs-only-pept-count**. Peptide counts for TAMs only proteins in all IP samples are provided. **Tumor-and-TAMs-motifs**. Peptides predicted in IP from tumor and tumor-associated monocytes/macrophages (TAMs) lysates that contained GH motifs are listed along with their size and location of GH motifs in mouse and human proteins. **Tumor-and-TAMs-pept-count**. Peptide counts for Tumor and TAMs proteins in all IP samples are provided. **GO_BP_FAT-GH-TAMs**. Functional annotation of all TAMs proteins (absent or decreased in controls) containing GH proteins using DAVID, results of GO_BP_FAT report. pH=pHis.

---

**Fig. S1:**
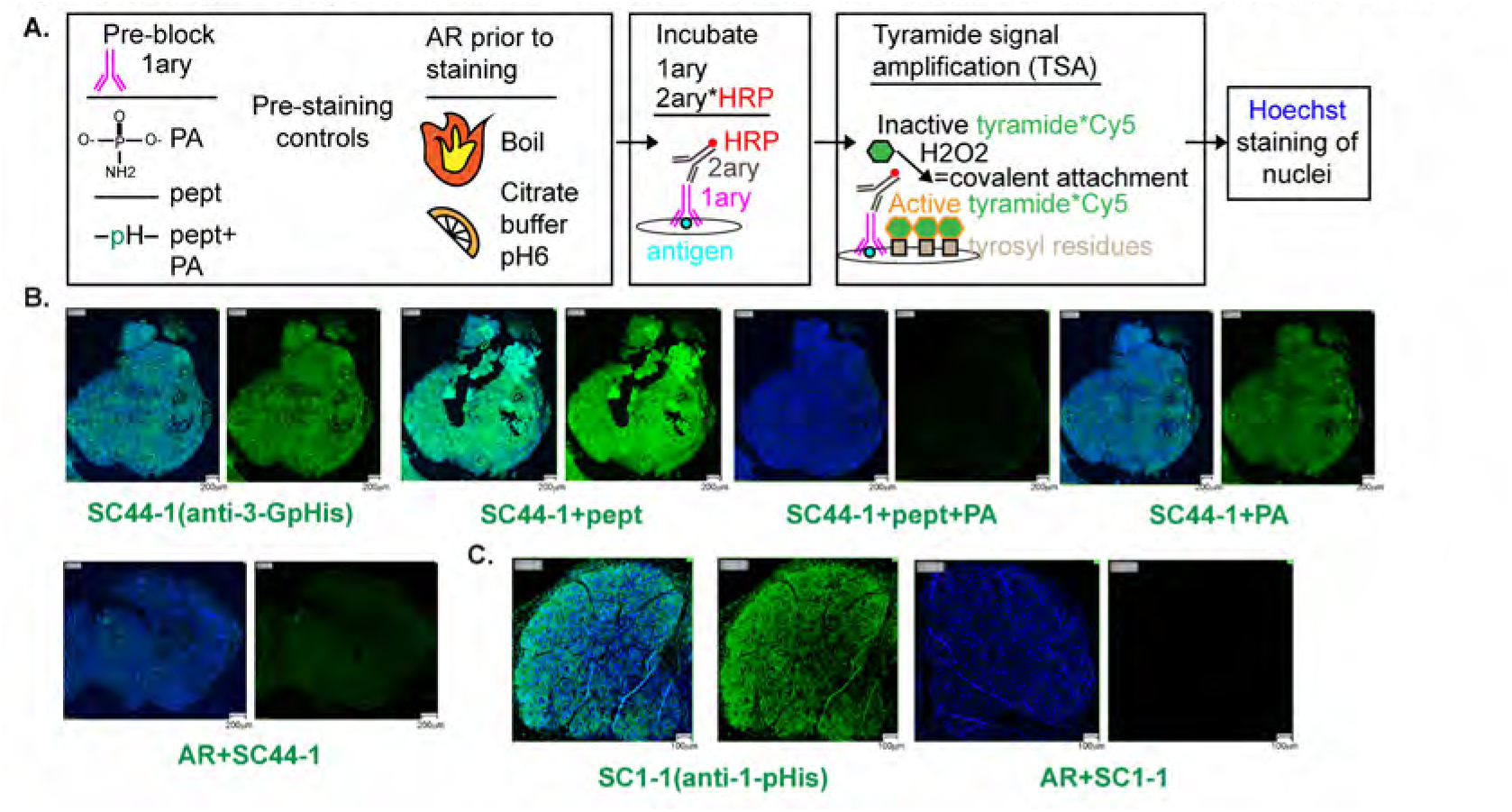
IHC-IF-pHis-TSA protocol enables detection of bona fide pHis signals in KP^*f/f*^*C* FFPE mouse PDAC tumor sections. (**a**), Schematic depicts the protocol used in this analysis. (**b-c**), Validation of antibody staining for the indicated anti-pHis antibody used with TSA Cy5 (green signals) reagent, Hoechst staining of nuclei in blue. PA=phosphoramidate, pept=unmodified peptide, pept+PA=phosphohistidine peptide, AR=antigen retrieval, 1ary=primary antibody, 2ary=secondary antibody. Scale bars are indicated at the lower right corner of images. PDAC=pancreatic ductal adenocarcinoma, pHis=phosphohistidine.

**Fig. S2.**
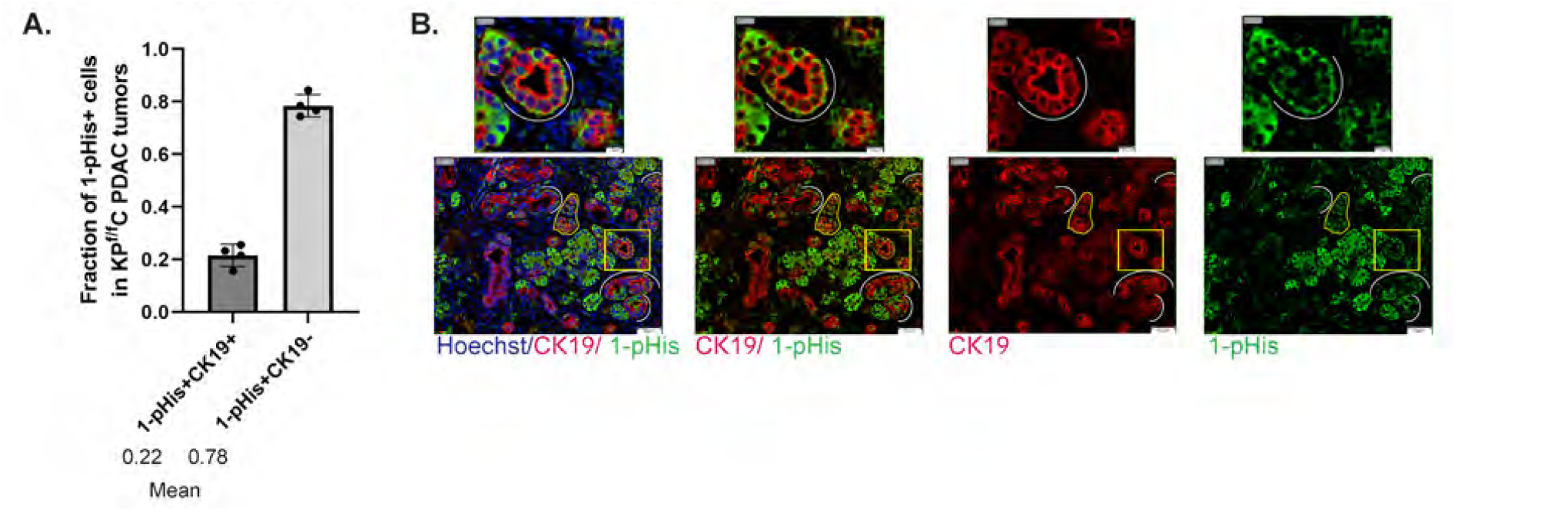
Expression of 1-pHis signals in mouse PDAC. IHC-IF-pHis-TSA protocol (Fig. S1) applied to FFPE mouse KP^f/f^C PDAC tumor sections, using the indicated primary antibodies with TSA Cy5 (green signals, 1-pHis) or TSA Cy3 (red signals, CK19) reagents, Hoechst staining of nuclei in blue. (**a**) Quantitation of the fraction of G-3-pHis+ cells co-staining with CK19+ cells for KP^f/f^C tumors, n=4. **(b**) A single slide-scanned image showing basolateral G-3-pH co-staining with CK19+ tumor ducts. The regions enclosed by the yellow boxes on the lower magnification images (bottom) are shown at higher magnification above. Yellow outline highlights an example of strong CK19 and 1-pHis co-staining. White lines show basolateral expression of 1-pHis on tumor ducts. Scale bars are indicated at the lower right corner of images. PDAC=pancreatic ductal adenocarcinoma, pHis=phosphohistidine.

**Fig. S3.**
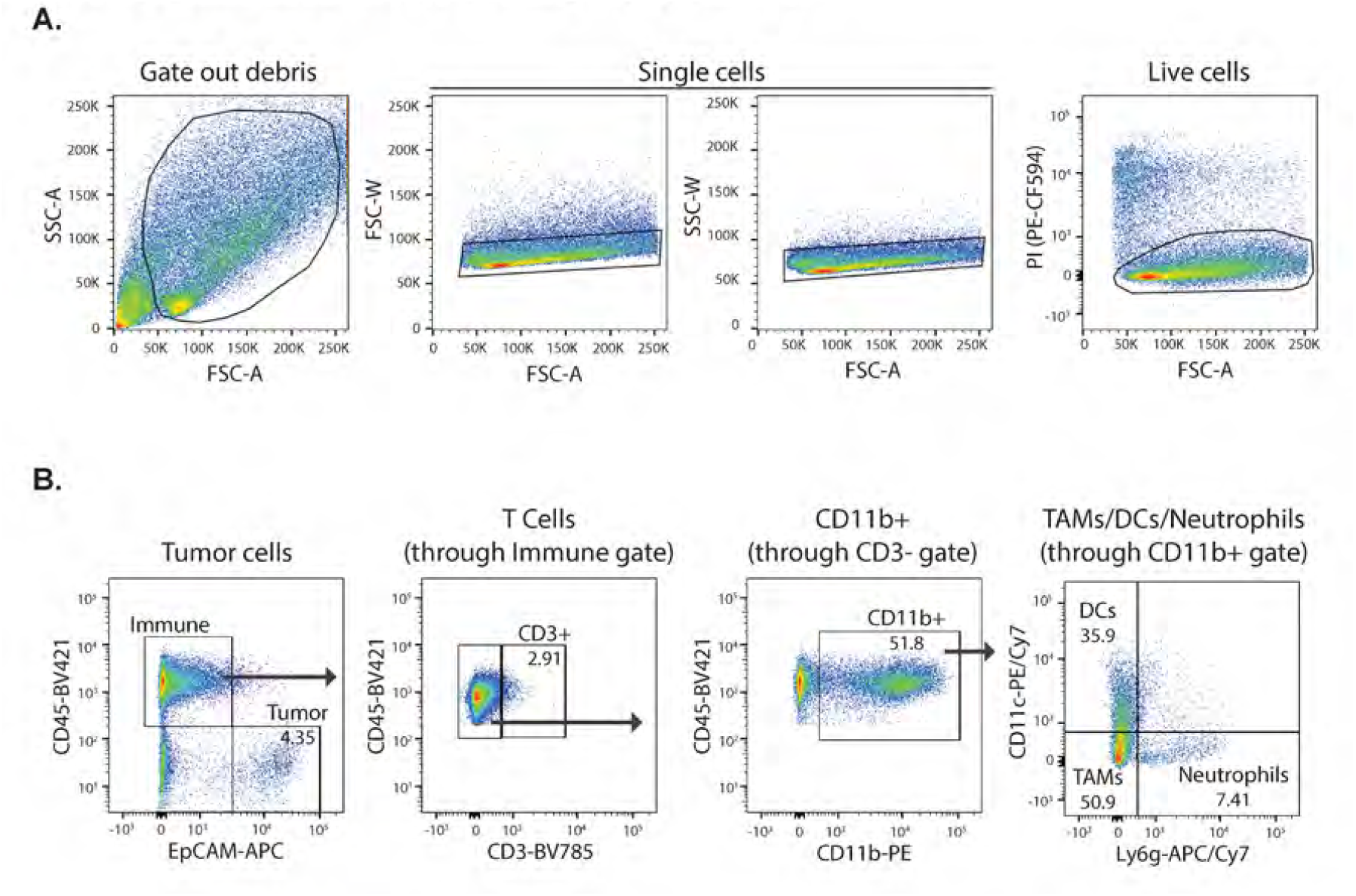
Scheme used for FACS sorting of populations. Examples of cell sorting plots for dissociated, advanced stage KP^f/f^C tumors. (**a**) Debris, single cell, live/dead cell gating (**b**) Gating for tumor, immune, T cell, dendritic cells (DCs), neutrophils, tumor-associated monocytes/macrophages (TAMs).

**Fig. S4.**
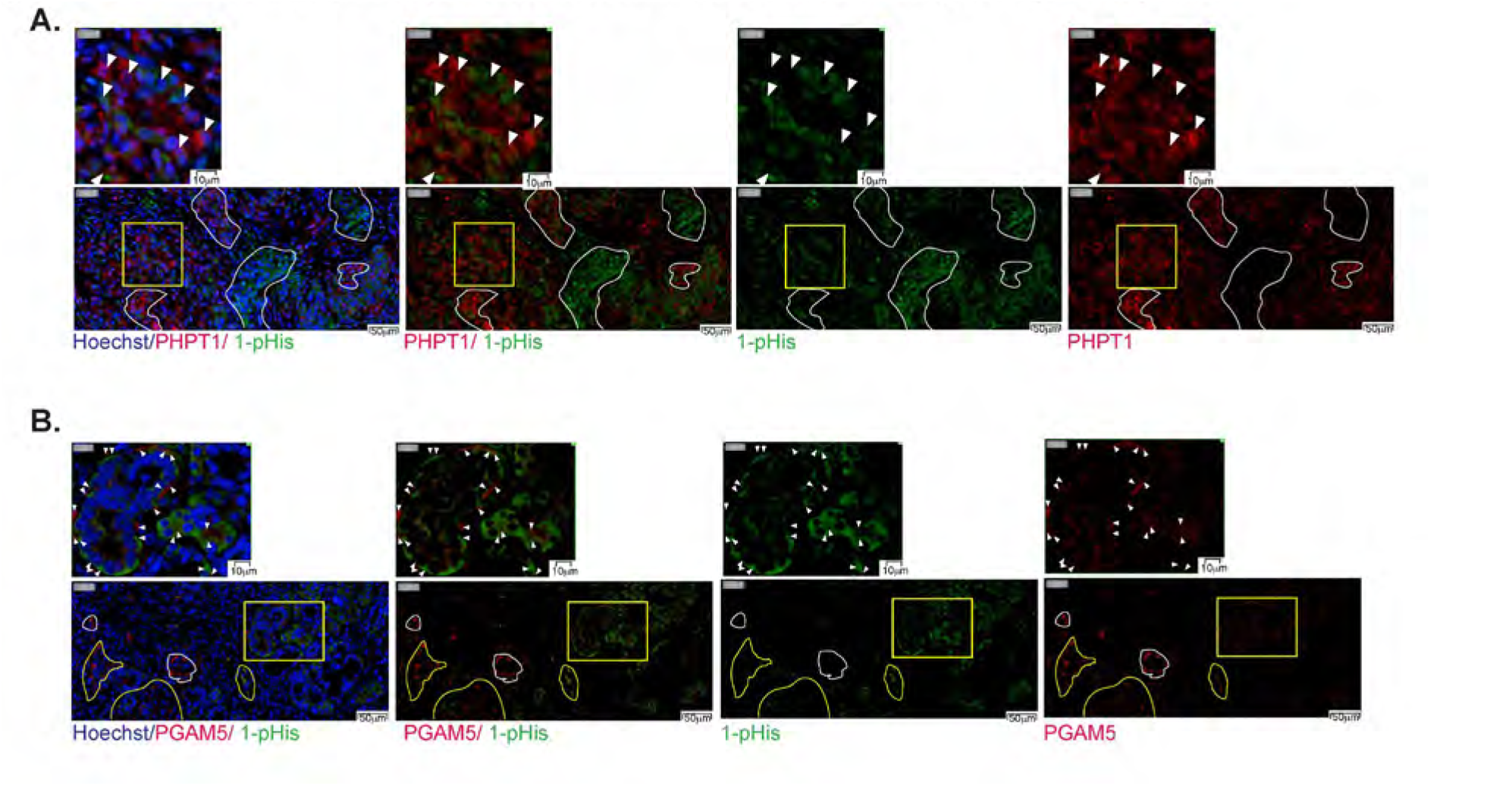
PHPT1 and PGAM5 signals exhibit an inverse relationship to 1-pHis signals. (**a-b)**, IHC-IF-pHis-TSA protocol (Fig S1) applied to FFPE KP^f/f^C mouse pancreatic tumor serial sections analyzed using the indicated primary antibodies with TSA Cy5 (green signals, 1-pHis) or TSA Cy3 (red signals, PHPT1 or PGAM5) reagents, Hoechst staining of nuclei in blue. Slide scanned images (20X, Olympus). The regions enclosed by the yellow boxes on the lower magnification images (bottom) are shown at higher magnification above. White outlines and arrowheads provide examples of red and green signals with an inverse relationship. Scale bars are indicated at the lower right corner of images. pHis=phosphohistidine.

**Fig. S5.**
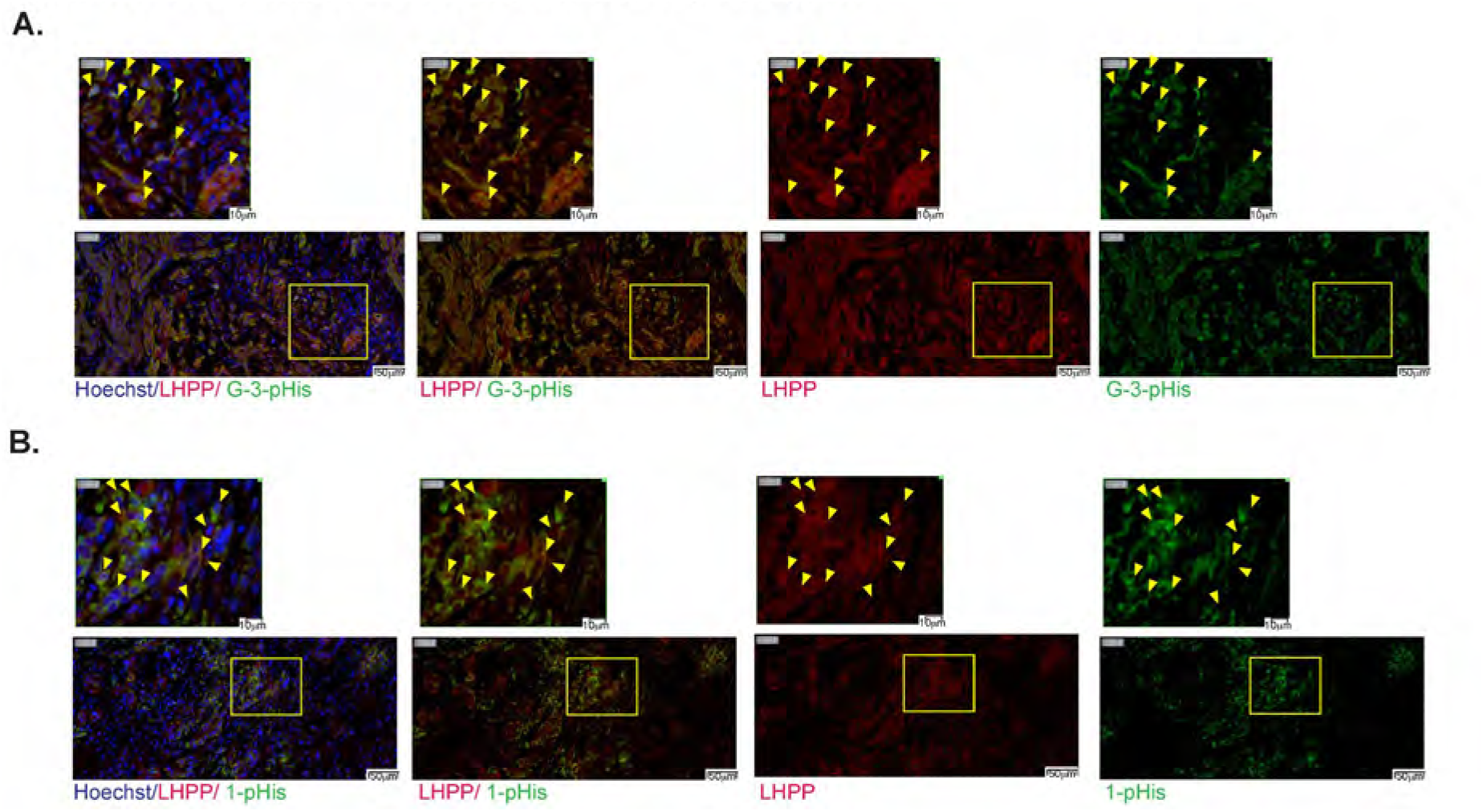
LHPP co-stains with G-3-pHis and 1-pHis signals. (**a-b)**, IHC-IF-pHis-TSA protocol (Fig S1) applied to FFPE KP^f/f^C mouse pancreatic tumor serial sections analyzed using the indicated primary antibodies with TSA Cy5 (green signals, G-3-pHis or 1-pHis) or TSA Cy3 (red signals, LHPP) reagents, Hoechst staining of nuclei in blue. Slide scanned images (20X, Olympus). The regions enclosed by the yellow boxes on the lower magnification images (bottom) are shown at higher magnification above. Yellow outlines and arrowheads provide examples of co-staining. Scale bars are indicated at the lower right corner of images. pHis=phosphohistidine.

**Fig. S6.**
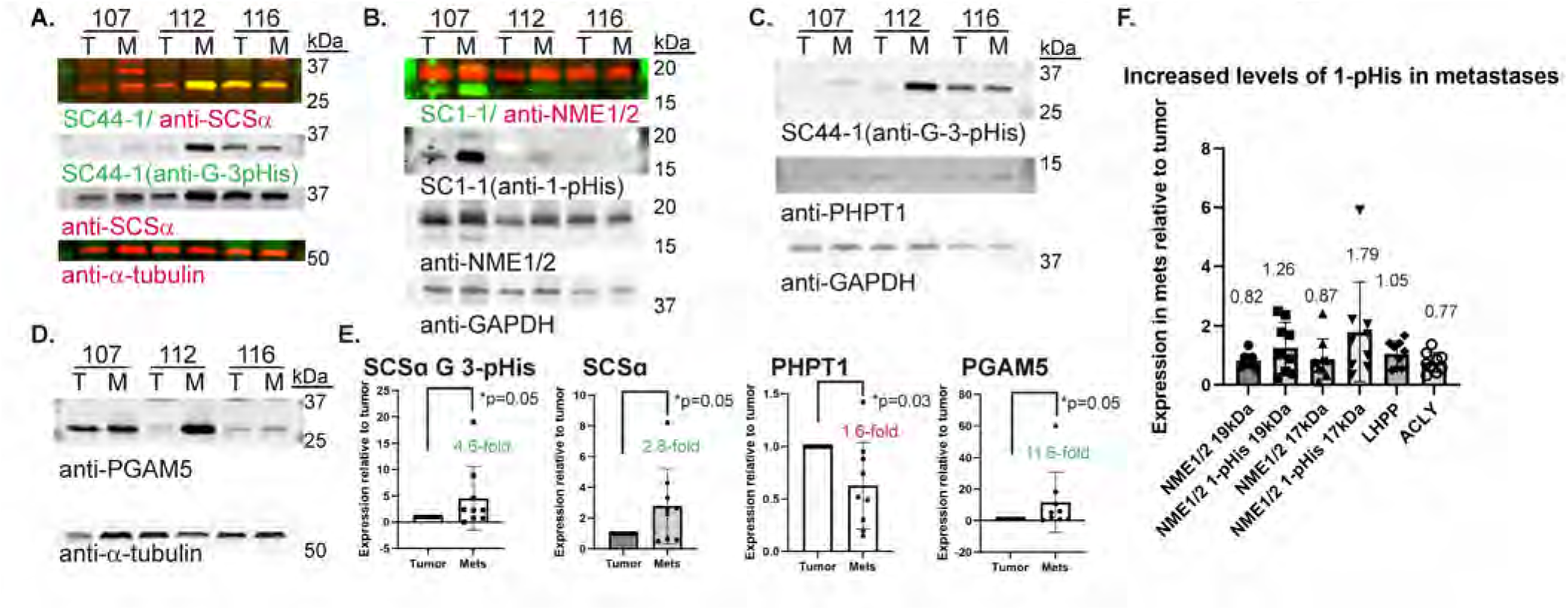
Altered expression of pHis and pHis regulators in human PDAC metastases: PGAM5 is a possible target. Immunoblotting (IB) and quantitation of tissue lysates from clinical paired human primary pancreatic tumor and metastases (n=9) (**a-d**) Representative IBs, each showing a single IB probed serially. Quantitation of α-tubulin or GAPDH-normalized expression in metastases relative to primary tumors for all IBs shown in graphs (**e**) (Wilcoxon) with average fold increase (green) and decrease (red) noted and in (**f**) with average expression in mets=metastases relative to tumor noted. pHis=phosphohistidine.

## Notes

### Competing Interest Statement

The authors have declared no competing interest.

